# Epithelial memory after respiratory viral infection results in long-lasting enhancement of antigen presentation

**DOI:** 10.1101/2024.07.26.605255

**Authors:** Piotr P. Janas, Wouter T’Jonck, Matthew O. Burgess, Caroline Chauché, Matthieu Vermeren, Christopher Lucas, Calum Bain, Robert Illingworth, Edward W. Roberts, Henry J. McSorley, Jürgen Schwarze

**Affiliations:** Centre for Inflammation Research, Institute for Regeneration and Repair, The University of Edinburgh, UK; School of Medicine, Medical Sciences and Nutrition, The University of Aberdeen, UK; CRUK Scotland Institute, Glasgow, UK; Division of Cell Signalling and Immunology, School of Life Sciences, University of Dundee, UK

## Abstract

**Background:** Viral lower respiratory tract infections (LRTIs) can reduce severity of subsequent LRTIs but have also been linked to respiratory allergy development and exacerbation. Here, we show that viral LRTI can imprint lung epithelial cells (LECs), leading to long-term phenotypic and functional changes in their response to subsequent challenges.

**Methods:** Mice were infected via intranasal administration of respiratory syncytial virus (RSV). After 28 days, LECs were isolated using cold dispase digestion followed by magnetic-activated cell sorting. Epigenetic changes were assessed using CUT&RUN, while transcriptional changes were evaluated using NanoString and qPCR. Flow cytometry was employed to measure cell surface MHC levels, antigen uptake and processing rates, and OT-I proliferation.

**Results:** We identified epigenetic and transcriptomic changes in murine LECs 28 days post respiratory syncytial virus (RSV) infection after recovery in genes associated with major histocompatibility complexes (MHC). Lasting upregulation of MHC-I and MHC-II was further increased following *in vivo* LPS stimulation. Importantly, MHC upregulation was associated with increased antigen uptake and processing, as well as increased antigen presentation to T cells.

**Conclusions:** Our data suggest that LRTI can induce long-term upregulation of antigen-presentation by LECs, thus facilitating local T cell responses to microbial antigens and allergens, potentially enhancing immunity or in susceptible hosts respiratory allergy.

## Introduction

Viral lower respiratory tract infections (LRTI) including rhinovirus (1), human metapneumovirus (2) and respiratory syncytial virus (RSV) (3,4) are linked to the development and exacerbation (5,6) of allergic airway disease but can also protect from subsequent respiratory infections (7,8). While multiple genetic, epigenetic and environmental factors predisposing to allergic airway disease development have been described (9,10), the underlying cellular and molecular mechanisms are still unclear. Historically, respiratory epithelial cells were viewed primarily as a relatively inert barrier at the interface of environment and host, however, in recent years their importance in orchestrating initial innate immune responses has been recognised (11). Given their relatively long lifespan (12), as well as reports that lung (13) and skin (14) epithelial cells can be phenotypically altered long-term, we hypothesised that viral LRTI can induce long-term changes in lung epithelial cells (LECs), that could contribute to responses to subsequent infection and allergen challenge. We used an established mouse model of RSV infection (15–17) to investigate long-lasting changes in LECs following viral clearance. Here, we reveal epigenetic and transcriptomic changes in genes primarily associated with major histocompatibility complex (MHC) and antigen processing that persist long-term after RSV infection. LECs also maintained elevated levels of both MHC-I and MHC-II with further enhancement after stimulation with lipopolysaccharide (LPS). These long-term changes were associated with increases in antigen uptake, processing and presentation to T cells.

## Materials and Methods

### RSV stock and immunoplaque assay

RSV A2 (kindly provided by Dr James Harker, Imperial College London) was expanded in Hep-2 cells as previously described (18). Once 10-20% drop in confluency was observed and syncytia were observed throughout, cells were sonicated, and virus was harvested. RSV titres were assessed using immunoplaque assay. Plaque forming unit (PFU/ml) was calculated based on the average plaque count and a dilution factor.

### Animals and animal procedures

Wild-type female BALB/c and C57BL/6J mice were procured from Charles River Laboratories. Mice were acclimatized for one week before experimental use. OT-I mice were imported from the CRUK Beatson Institute, Glasgow. Both male and female OT-I mice ranging from two to six months old were used for OT-I cell harvest. Mice were housed in individually ventilated cages. Animal work was carried out under the regulations of the Animals (Scientific Procedures) Act 1986. All procedures were approved by the University of Edinburgh Animal Welfare and Ethical Review Board and performed under UK Home Office licenses with institutional oversight performed by qualified veterinarians. UK Home Office project license to JS, number PP4544912. ARRIVE 2.0 guidelines were followed were applicable.

6-week-old BALB/c or C57BL/6J mice were administered 50µl of 5x10^6^/ml RSV A2 intranasally under light anaesthesia (inhaled isoflurane). Alternatively, mice were administered 50µl of PBS (w/o Ca^2+^ and Mg^2+^) or UV-irradiated RSV (using SPECTROLINKER XL-1500 at 2J/cm^2^). After 28 days mice were either processed as described below or intranasally administered 50µl PBS (w/o Ca^2+^ and Mg^2+^) or 10µg of LPS (Sigma-Aldrich, USA, E. coli O111:b4) in 50µl PBS for either 6h or 24h before further processing as described below.

### Murine lung harvest and processing

Murine lungs were harvested and processed as previously described (19). In brief, lungs were inflated with ice-cold 1.5-2ml enzyme mix (DMEM/F12 (Gibco, USA) + 100U/ml Pen/Strep (Gibco, USA) + 2mg/ml Dispase II (Sigma-Aldrich, USA) + 0.1mg/ml DNaseI (Sigma-Aldrich, USA)). Lungs were then incubated at 4-6°C for 20h. After incubation, lungs were passed through 70µm and 30µm strainers and red blood cells (RBC) were lysed for 2min using ACK RBC lysis buffer (Gibco, USA). Cells were resuspended in MACS buffer (PBS w/o Mg2^+^ and Ca2^+^ + 0.5% bovine serum albumin (BSA) (Sigma-Aldrich, USA) + 2mM EDTA (Gibco, USA) + 100U/ml Pen/Strep (Gibco, USA)). The cell suspension was then either resuspended in the appropriate media for *in vitro* culture (discussed below) or incubated with 5µl/1ml/lung anti-mouse CD16/32 antibody (BioLegend, USA) for 30min at 4°C before further processing.

### Isolation of lung epithelial cells

LECs were MACS-isolated as previously described (19). In brief, up to 10^7^ cells were incubated with 5µl anti-CD31 microbeads and 10µl of anti-CD45 microbeads (Miltenyi Biotec, Germany), followed by LS column magnetic isolation. Flowthrough cells were incubated with 15µl of anti-EpCAM microbeads (Miltenyi Biotec, Germany) followed by magnetic isolation with MS columns. CD45-CD31-EpCAM+ LECs were then flushed out from MS columns with a plunger, and their purity (>97%) was assessed using flow cytometry (Figure S1).

### Cleavage Under Targets and Release Using Nuclease (CUT&RUN)

2x10^5^ of MACS-sorted murine LECs from a pool of three mice were used per single CUT&RUN reaction. Unless specified otherwise all sample incubations were carried out at 4°C. Cells were washed twice using PBS with 600 x g centrifugation steps at 4°C in-between each wash. Cells were then incubated in 10% Triton X-100 (Sigma-Aldrich, USA) for 10min to isolate nuclei. Nuclei were pelleted following 1300 x g centrifugation at 4°C for 5min and resuspended in wash buffer. Following nuclear isolation, a CUT&RUN protocol (20) was followed. In brief, nuclei were bound to BioMag Plus Concavalin A beads (Bangs Laboratories, USA) and resuspended in antibody buffer containing either anti-H3K27ac (1:100, Active Motif, USA), anti-H3K4me3 (1:100, Sigma-Aldrich, USA) or normal rabbit IgG (1:100, Cell Signalling Technology, USA). Following overnight incubation, nuclei were washed thrice with wash buffer and incubated with protein A/G-MNase (VIB Protein Core, kindly provided by Prof. Martin Guilliams) for 1 hour while rotating. Chromatin digestion and release was performed using the high Ca^2+^/low salt method followed by phenol/chloroform extraction. DNA was stored at –20 °C until further processing. KAPA Prep Kit (Roche, Switzerland) and KAPA UDI adapter kit (Roche, Switzerland) were used for library preparations per manufacturer’s instructions. In brief, end repair 5’ phosphorylation (20°C, 30min), dA-tailing (58°C, 45min) and adapter ligation (20°C, 60min) were followed by un-ligated adapters removal with 1.1x volume of AMPure XP beads (Beckman Coulter, USA). Libraries were amplified over 12 cycles in thermocycler using KAPA HiFi HotStart ReadyMix and 5µM Pre-LM-PCR Oligo 1&2 (Roche, Switzerland). Following amplification another 3-step adapter removal with AMPure XP beads was carried out. Samples were quality controlled (Figure S2) using LabChip GX DNA High Sensitivity (Caliper Life Sciences, USA) according to manufacturer’s instructions by biomolecular core staff at Institute for Regeneration and Repair (IRR), University of Edinburgh.

Libraries were then submitted to Azenta Life Sciences for next generation sequencing using 2x150bp configuration at 5x10^6^ reads per sample. Fastq sequencing files were uploaded to Galaxy platform (21) for QC using FastQC (22) and sequence trimming using Trimmomatic (23), followed by sequence alignment to mm10 mouse reference genome using Bowtie2 (24). Next, bamCoverage (25) tool was used to generate bin size 5 read coverage .bam files, followed by peak calling using Sparse Enrichment Analysis for CUT&RUN (SEACR) (26). Bowtie2 and SEACR files were exported and analysed further using R Studio R4.3.1. DiffBind package (27) (release. 3.14) to identify differentially bound sites (DBS). Duplicate reads were not removed, while problematic regions were removed using ENCODE Blacklist database (28). Statistical discovery of significant DBS was carried out using DESeq2 (29). Gene annotation and genomic annotation were carried out using ChIPSeeker package (30) (release 1.4), while gene ontology (GO) analysis was carried out using ClusterProfiler (31) (release 4.12).

### RNA extraction

RNA was extracted as previously described (19). In brief, 100µl of bromochloropropane (Sigma-Aldrich, USA) was added to cells treated with TRizol (Life Technologies, USA), followed by 16000 x g at 4°C, 20min centrifugation and RNA precipitation using isopropanol. RNA pellets were then washed three times using 70% EtOH. After determining RNA concentration samples were normalised using RNase-free H_2_O, followed by treatment with DNase I (QIAGEN, Germany) as per manufacturer’s instructions to remove genomic DNA.

### NanoString

All RNA samples were determined to be of high quality as determined by RNA 6000 Pico Assay (Agilent, USA) >80% DV200. RNA from two mice was pooled to generate a single sample. Each sample was normalised to 20ng/µl, and 8µl/sample of RNA was submitted to Host Tumour Profiling Unit Microarray Services for profiling with Nanostring nCounter Mouse Immunology Panel. NanoString nCounter protocol was followed according to the manufacturer’s instructions. NanoString data was analysed using “NanoTube” package (32) (release 3.18) in RStudio R4.3.1 as previously described (33). The bgPval was set to 0.01 and the number of unwanted factors (k) was set to 3, while the remaining normalization parameters were set to default.

### qPCR

qPCRs were performed as previously described (19). Each sample was normalized to appropriate endogenous control within the reaction and log2FC was calculated using ΔΔCt method. Appropriate endogenous controls were selected using “NormFinder” package (34) (release 0.1.2). *Oaz1* and *Gapdh* were identified as the most stably expressed endogenous controls between experimental groups (**Table S1**). *Oaz1* was used for low expressed targets, while *Gapdh* was used for highly expressed targets. qPCR data is presented as log2 fold change relative to PBS sample.

### Confocal immunofluorescent microscopy

Murine lungs were inflated using 0.8ml of 1:1 PBS/OCT mixture and placed in antigenfix (Diapath, Italy) for 45min at RT. Lungs were then washed twice using PBS and placed in 34% sucrose solution for 24h at 4°C. Lungs were snap frozen in OCT and sliced (7µm sections). Following rehydration, sections were incubated in 100µl of blocking buffer (Tween-20, 2% BSA, 5% FBS (LabTech, UK), 2% mouse serum (Thermo Fisher Scientific, USA) and 5% normal goat serum (Thermo Fisher Scientific, USA) in PBS) for 1.5h at RT in humidity chamber. Fluorochrome-conjugated antibodies (1:400 anti-EpCAM-AF594, 1:200 anti-MHC-I-AF647 and 1:400 anti-MHC-II-AF488 (BioLegend, USA)) were diluted in blocking buffer and 100µl of staining mix was placed on each section for 1h at RT. Slides were imaged using a Leica SP8 confocal microscope using 20x/0.75 objective without immersion or 40x/1.3 objective with oil immersion objectives. A minimum of 3 fields of view were captured per each section. Images were imported into Fiji and nuclear StarDist (35) segmentation refined by EpCAM expression stratifying to the airway (EpCAM^high^) and alveolar (EpCAM^low^) regions was carried out (**Figure S3**). The mean fluorescence intensity of MHC-I (AF647) and MHC-II (AF488) channels was measured per each section. Segmentation was visually assessed, and each section with incorrect segmentation was eliminated from further analysis (in total 7 sections were removed out of the 84 sections imaged).

### Flow cytometry

Staining for flow cytometry analysis was performed as previously described (19). In brief, following incubation with anti-CD16/32 antibody cells were stained using a LIVE/DEAD fixable near-IR stain (Invitrogen, USA), followed by staining with appropriate antibody cocktail (**Table 2**). For intracellular TLR4 staining, the Bioscience Foxp3/Transcription Factor Staining Buffer Set (Invitrogen) was used according to the manufacturer’s protocol. Samples were unmixed (with multiple autofluorescence extraction) using Cytek Aurora with Cytek SpectroFlo 3.3 and analysed using De Novo Software FCSexpress 7. The gating strategy involved debris exclusion (side scatter/forward scatter – SSC-H/FSC-H), followed by singlets selection (FSC-H/FSC-A), RBC exclusion (SSC-H/SSC-B-H) and dead cell exclusion (LIVE/DEAD Fixable Near-IR/FSC-H), followed by further experiment-specific gating (**Figure S4, Figure S5**). Flow cytometry data is presented as fold change relative to average median fluorescence intensity (MFI) of PBS samples. We validated our EpCAM stratification strategy to airway and alveolar LECs using CD49f (airway epithelial marker (36)), MHC-II (pneumocyte marker (37)), and CD24 (conducting airways marker (38)) (**Figure S6**).

**Table 1.** TaqMan qPCR probe details.

| Target | Species | Probe | Assay ID | Catalogue number | Endogenous control |
| --- | --- | --- | --- | --- | --- |
| *Gapdh* | Mouse | VIC-MGB | Mm99999915_g1 | 4448489 | NA |
| *Oaz1* | Mouse | VIC-MGB | Mm07307469_g1 | 4448489 | NA |
| *H2-DMb2* | Mouse | FAM-MGB | Mm00783707_s1 | 4331182 | *Oaz1* |
| *B2m* | Mouse | FAM-MGB | Mm00437762_m1 | 4331182 | *Gapdh* |
| *H2-Ab1* | Mouse | FAM-MGB | Mm00439216_m1 | 4331182 | *Gapdh* |
| *H2-Eb1* | Mouse | FAM-MGB | Mm00439221_m1 | 4331182 | *Oaz1* |
| *Casp4* | Mouse | FAM-MGB | Mm00432304_m1 | 4331182 | *Oaz1* |
| *Psmb9* | Mouse | FAM-MGB | Mm00479004_m1 | 4331182 | *Oaz1* |
| *Tap1* | Mouse | FAM-MGB | Mm00443188_m1 | 4331182 | *Oaz1* |

**Table 2.**
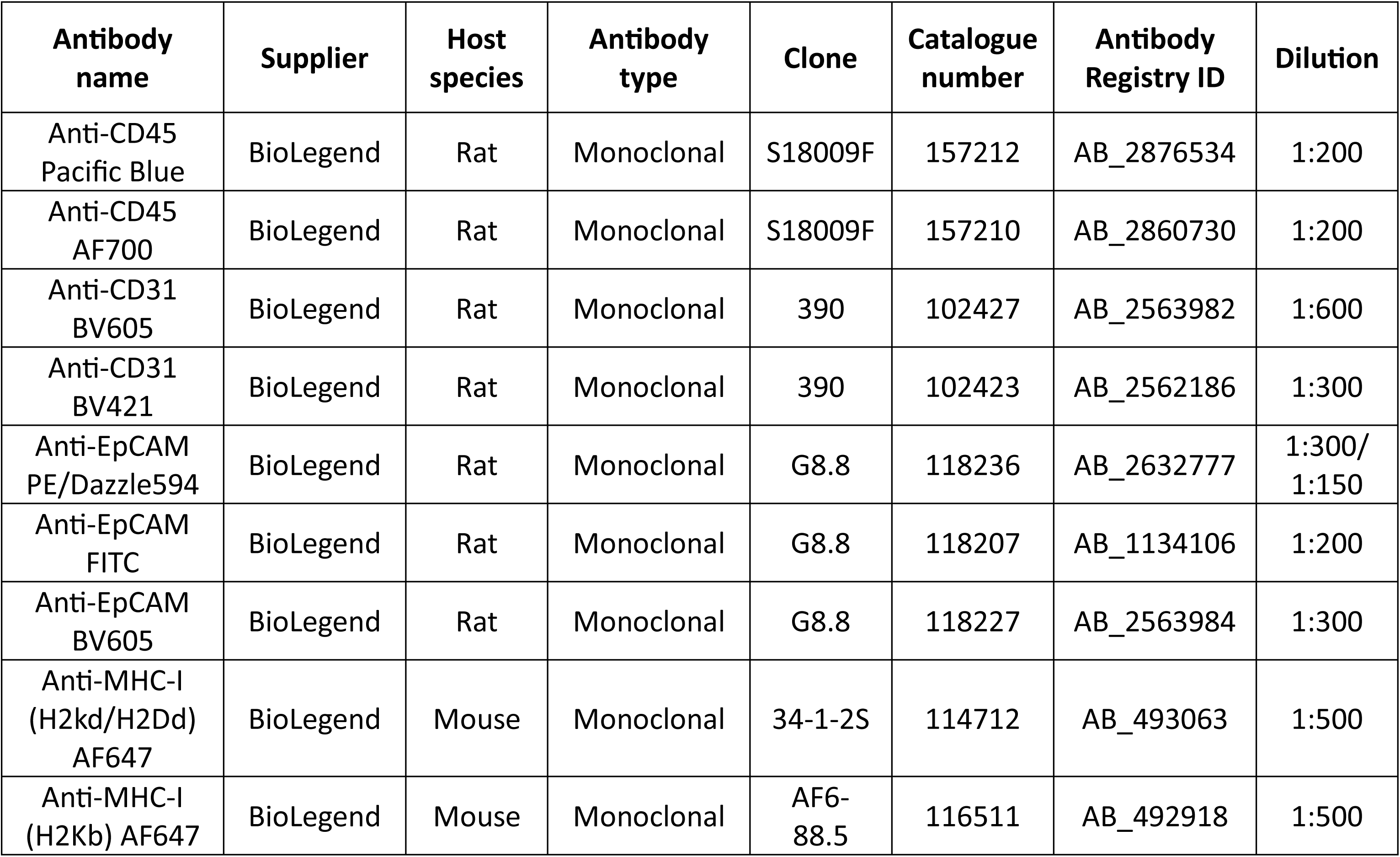
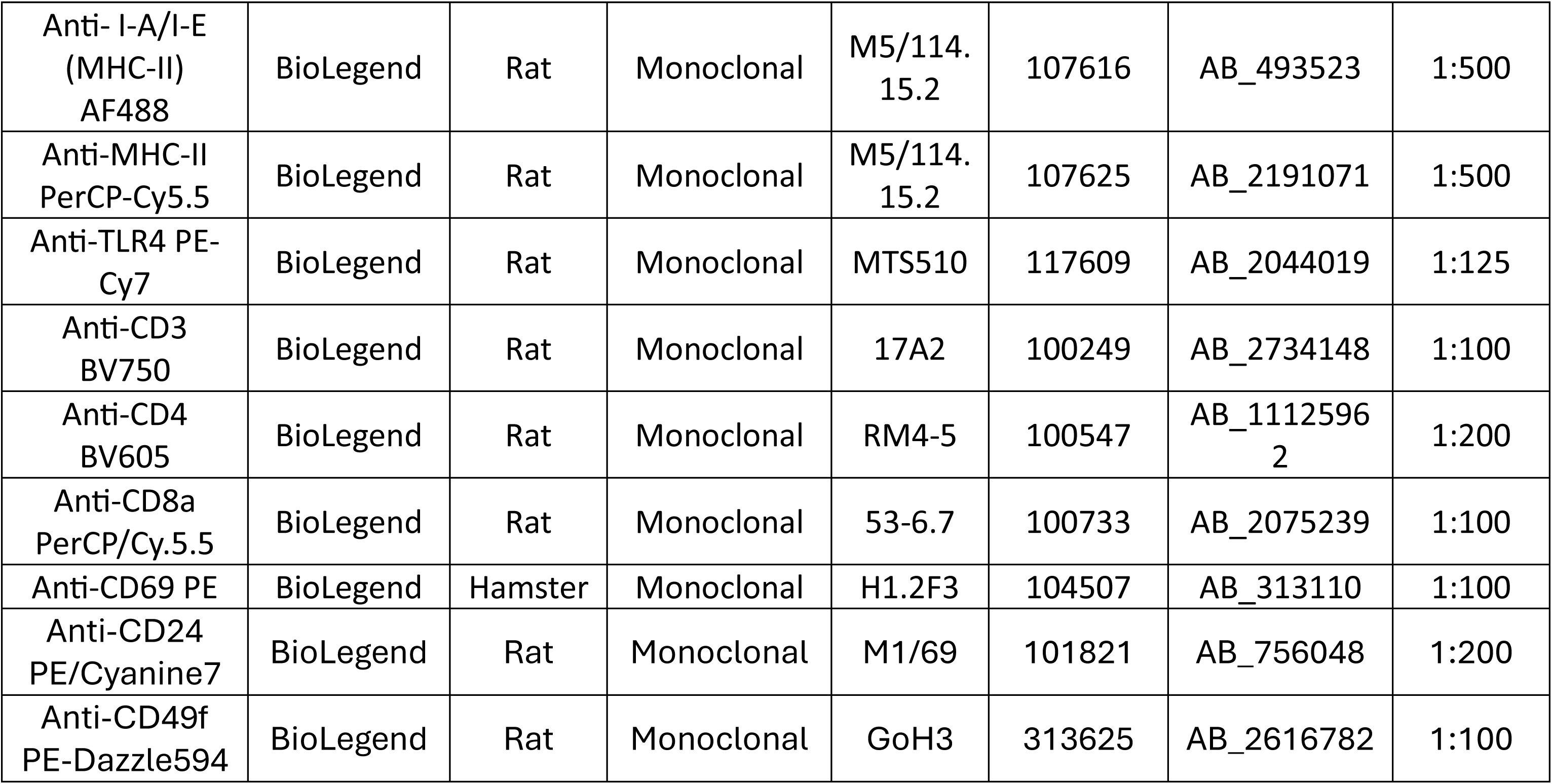
Flow cytometry antibodies.

### Antigen uptake and processing assays

Following lung harvest and LEC isolation, 1x10^6^ live cells (as determined by trypan blue) were resuspended in 180µl of pre-warmed serum-free PromoCell Airway Epithelial Growth Medium. Following addition of 20µl of OVA-AF647 (Invitrogen, USA) and DQ-OVA (Invitrogen, USA)) at a final concentration of 10µg/ml cells were incubated at 37°C 5% CO2 for 1h or 4h. All samples were collected simultaneously for further antibody staining and flow cytometric analysis (as described above).

### T cell proliferation assays

Isolated LECs were placed in complete T-cell media (RPMI + 10% FBS + 1% L-Glu + 100U/ml Pen/Strep + 1% MEM NEAA + 1% Na-Pyruvate + 0.1% 2-mercaptoethanol) (Gibco, USA) with PBS or LPS (1µg/ml) and SIINFEKL peptide (InvivoGen, USA) at 25µg/ml for 4h at 37°C 5% CO_2_. Meanwhile, inguinal, mesenteric, brachial, axillary and superficial cervical lymph nodes (LNs) were harvested from OT-I mice into cold complete T-cell media. Lymph nodes were mashed and then processed using the lung harvest and processing protocol as described above. CD8 T-cell MojoSort (BioLegend, USA) protocol was followed for negative labelling of CD8 T-cells. Up to 5x10^7^ labelled cells were placed in the Miltenyi LS column on the OctaMACS magnet. Flowthrough was collected and cells were stained with 1:5000 CFSE (Invitrogen, USA) according to manufacturer’s instructions. 1x10^5^ OT-I cells were added per 2x10^4^ LECs in 96 round-bottom well plate. Cells were co-cultured in 200µl complete T-cell media for 72h at 37°C 5% CO_2_, followed by antibody staining and flow cytometric analysis as described above (**Figure S5**). Proliferation index was calculated as the average number of cells that an initial cell became.

### Data presentation and statistical analysis

Data is represented as violin, split violin, scatter or dot plots generated using “ggplot2” package (39) (release 3.18). “webr” package (release 0.1.5) was used to create donut charts and “UpSetR” package (release 1.4.0) (40) was used to create an UpSet plot. Dark shaded areas represent the actual distribution of data, while lightly shaded areas represent the predicted probability density. Solid lines represent the median, while dashed lines represent 0, 25th, 75th and 100th percentile quartiles. R studio “rstatix” package (41) (release 0.7.2) was used for statistical analysis. The normality of the data was assessed using the Shapiro-Wilk test. For single-variable analysis, a One-Way ANOVA was employed, followed by Holm post-hoc test for multiple comparisons (where more than two groups were present). For dual-variable analysis of multiple groups, a Two-Way ANOVA was employed, followed by Holm post-hoc test for multiple comparisons. Statistical comparisons were added to plots using “ggpubr” package (42) (release 0.6.0).

## Results

### RSV infection results in long-lasting epigenetic and transcriptional changes in LECs

To test our hypothesis that RSV infection results in long-lasting changes in LECs, we employed a murine model of RSV infection and investigated epigenetic and transcriptomic changes long after viral clearance. Specifically, we focussed on 28dpRSV given the body of data showing that RSV is cleared within seven days (43–45) (**Figure 1A**). First, to interrogate potential long-lasting epigenetic changes, we performed CUT&RUN of histone 3 lysine 4 trimethylation (H3K4me3) and histone 3 lysine 27 acetylation (H3K27ac); both histone modifications associated with transcriptional activation when present within the regulatory region of a gene (46–48). We discovered 197 differentially bound sites (DBS) across both histone modifications in RSV LECs 28dpRSV compared with controls (**Figure 1B-C, Table S2).** Importantly, the vast majority of identified DBS were associated with regulatory elements of corresponding genes; more than 72% of H3K27ac and 90% of H3K4me3 DBS were identified in promoter regions, suggesting that observed changes in histone modification enrichment may be functional (**Figure 1D**). Although both enriched and depleted DBS were identified, only enriched DBS exhibited strong statistical significance and magnitude of change (**Figure 1E**). Gene ontology term analysis identified viral and interferon responses, and several pathways related to major histocompatibility complexes (MHC) and antigen processing (**Figure 1F**).

**Figure 1.**
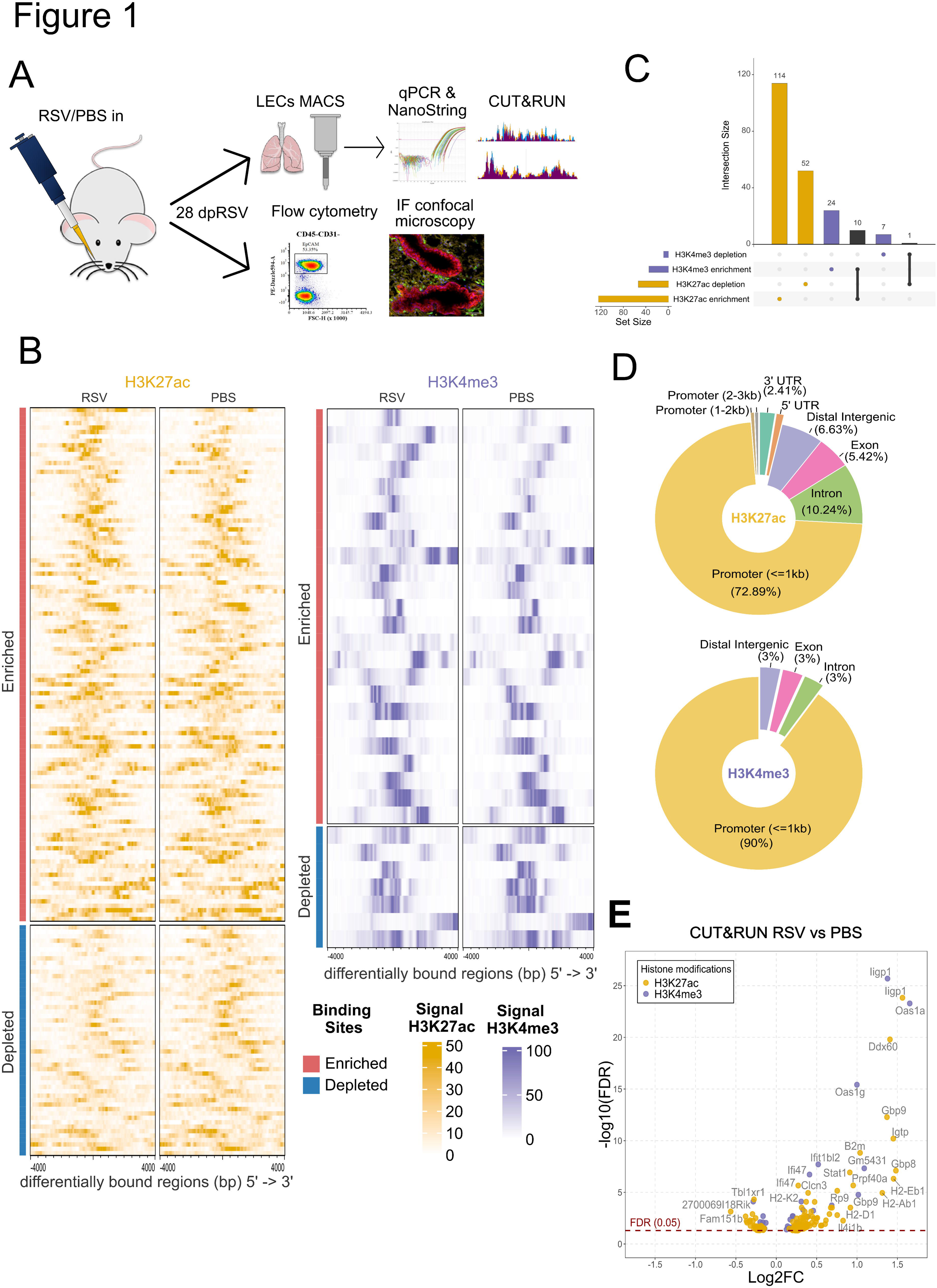
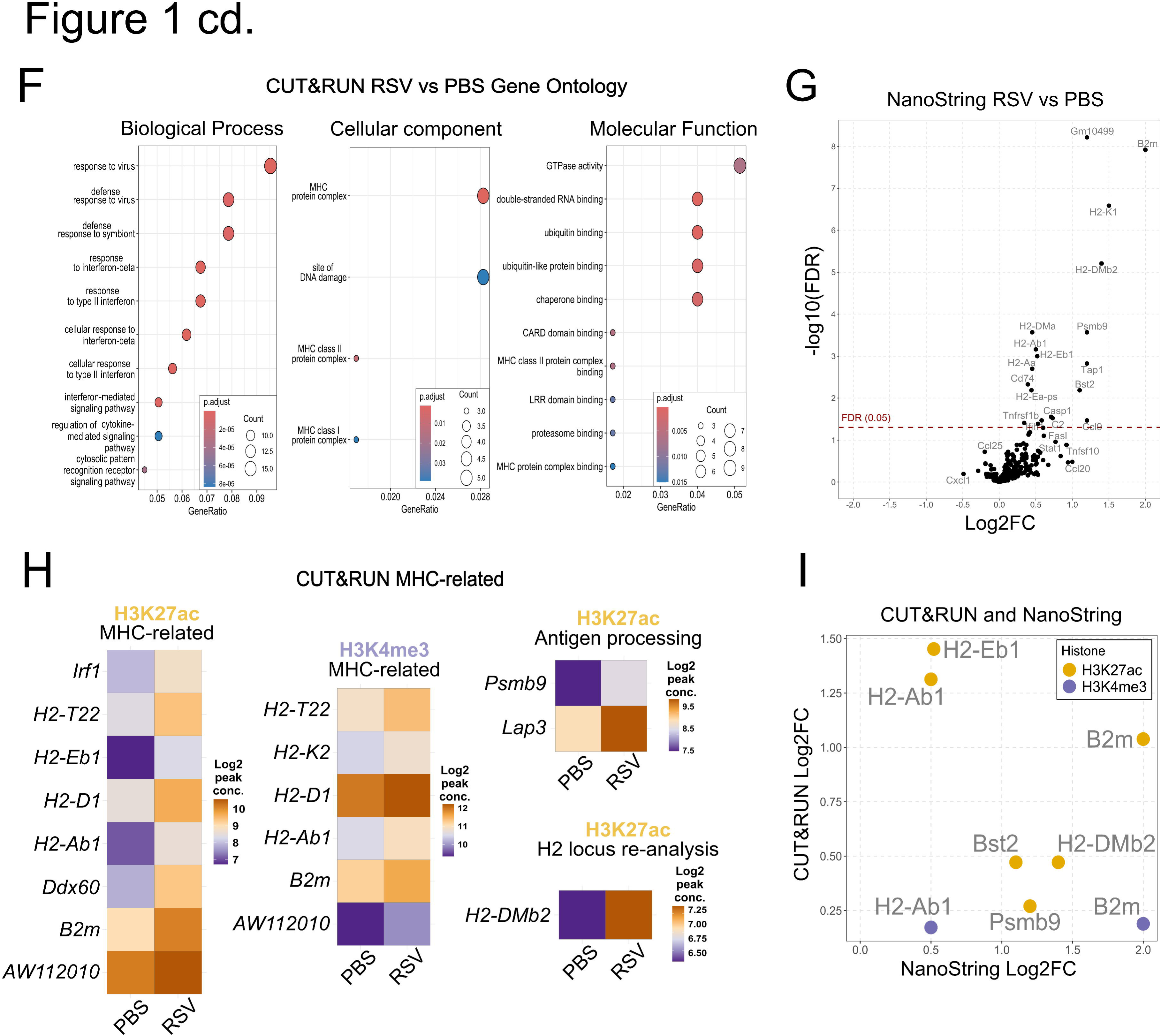
RSV cause long-term epigenetic changes in LECs. (**A**) Experimental design. 6-week-old female BALB/c mice were administered RSV or PBS intranasally. After 28 days mice were culled and lung epithelial cells were MACS-sorted for NanoString or CUT&RUN protocols, or lungs were harvested for confocal immunofluorescence microscopy or flow cytometry analysis. (**B**) CUT&RUN peak profile heatmaps of significant differentially bound H3K4me3 and H3K27ac regions – 8kbp window. (**C**) UpSet plot of identified enriched and depleted DBS for H3K4me3 and H3K27ac (**D**) Donut charts of annotated genomic distribution of identified DBS for H3K4me3 and H2K27ac. (**E**) Volcano plot of genes associated with H3K4me3 and H3K27ac that were differentially bound in RSV group as compared to PBS group (FDR<0.05). X axis shows a log2 fold-change in enrichment/depletion for each histone modification, while y axis shows a - log10 FDR values. (**F**) Dot plot graph of gene ontology (GO) term enrichment for combined H3K4me3 and H3K27ac. GO terms were stratified based on one of three sub-ontologies: biological process (BP), cellular component (CC) and molecular function (MF). Surface area of each dot corresponds to the number of genes assigned to each term, while colour levels correspond to statistical significance of discovered term. Gene Ratio on x axis is a ratio of genes identified per GO term over the total number of genes assigned to that term. (**G**) Volcano plot of differentially expressed LEC genes 28dpRSV identified by NanoString nCounter mouse Immunology panel in RSV group as compared to PBS group. X axis shows a log2 fold-change in expression, while y axis shows -log10 FDR values. (**H**) Heatmaps of relevant MHC or antigen processing-related genes identified by CUT&RUN that were enriched in RSV group. Murine H2 locus was re-analysed, resulting in identification of one additional, previously unreported, target – *H2-DMb2*. (**I**) Scatter plot of targets identified by both CUT&RUN and NanoString 28dpRSV. Y axis shows log2 fold-change in histone enrichment, and corresponding log2 fold-change in gene expression on x axis as identified by NanoString. CUT&RUN n=4 per group. NanoString n=6 per group. Each CUT&RUN sample is a pool of three mice, and each NanoString sample is a pool of two mice. Each experiment was repeated once.

Next, to determine the impact of RSV infection on the transcriptional profile of LECs, we used a NanoString Immunology panel. We discovered nineteen differentially expressed genes; strikingly eleven of those were associated with both classes of MHC and antigen processing (**Figure 1G**). This led us to reassess the CUT&RUN data in the context of MHC-related genes. Of the 197 significant DBS, eight H3K27ac and six H3K4me3 were associated with MHC biology, while two H3K27ac DBS were associated with antigen processing (**Figure 1H**). Additionally, CUT&RUN analysis specifically within the H2 locus, where the majority of MHC-related genes are located (49) revealed a single novel enriched H3K27ac DBS in the promoter of *H2-DMb2*. There was also overlap between long-term epigenetic and transcriptomic changes. Six genes identified as differentially enriched by CUT&RUN were also differentially expressed in NanoString data (**Figure 1I**). Five of the genes are associated with MHC and antigen processing, while one gene, *Bst2*, is thought to be involved in antiviral responses (50). Taken together, these data demonstrate that RSV infection results in long-term transcriptional and epigenetic effects on LECs.

### RSV infection alters expression of MHC and associated apparatus by LECs

Following the identification of several differentially expressed and histone-modified MHC-related genes, we validated upregulation of several representative genes by qPCR (**Figure 2A**). We further used confocal microscopy to validate MHC upregulation at the protein level (**Figure 2B-C**), finding higher levels of both MHC-I and MHC-II in the airway and alveolar compartments 28dpRSV. Finally, MHC expression was assessed at single cell level by flow cytometry, which confirmed that CD45-CD31-EpCAM^high^ airway LECs and CD45-CD31-EpCAM^low^ alveolar LECs (**Figure S6**) retained high expression of both MHC-I and MHC-II at 28dpRSV compared with PBS and UV-RSV controls (**Figure 2D**), indicating that persistent MHC upregulation requires RSV infection and not just RSV-antigen. Considering this, we investigated whether the magnitude of MHC expression correlates with infection severity by correlating weight change at 6dpRSV (peak infection severity (45,51)) with MHC expression increases 28dpRSV (**Figure S7**). Using Pearson correlation coefficient, we found that expression increases of MHC-I in both lung compartments and of MHC-II in airways strongly correlate with infection severity. Thus, these data show that increased MHC expression is maintained long after RSV infection is cleared and is associated with an altered epigenetic landscape.

**Figure 2.**
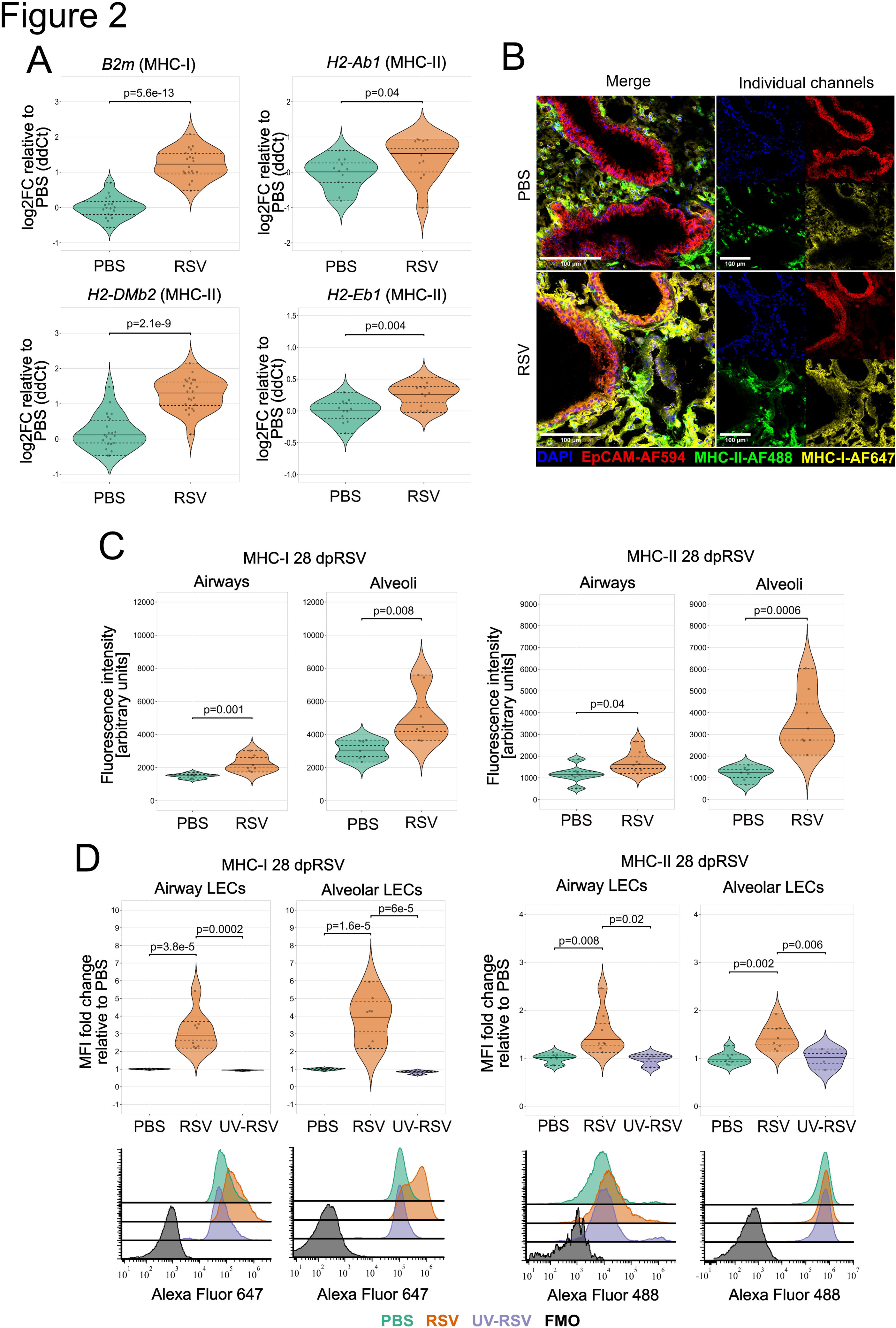
RSV infection results in long-term upregulation of MHC-I and MHC-II on LECs. 6-week-old female BALB/c mice were administered RSV, PBS or UV-RSV intranasally. 28dpRSV LECs were MACS-sorted for qPCR, or lungs were processed for flow cytometry or confocal IF microscopy. (**A**) Duplex TaqMan qPCR validation of selected targets identified by CUT&RUN and NanoString. qPCRs were performed on MACS-sorted LECs. *Oaz1* was used as endogenous control for *H2-DMb2* and *H2-Eb1* while *Gapdh* was used as endogenous control for *B2m* and *H2-Ab1*. Data presented as log2FC relative to PBS. N=12-24. (**B**) Representative confocal IF micrographs of MHC distribution in sucrose-dehydrated 7µm-thick OCT-embedded murine lung 28dpRSV. 40x magnification, white scale bar – 100µm. Blue DAPI – nuclei, red – EpCAM-AF594, green – MHC-II-AF488, yellow – MHC-I-AF647. (**C**) Quantification of absolute fluorescence intensity (MHC-I and MHC-II) in 20x magnification micrographs in airways and alveoli based on EpCAM expression via automated StarDist segmentation. N=7-8. (**D**) Relative flow cytometry quantification of MHC-I and MHC-II expression 28dpRSV stratified based on EpCAM expression which corresponds to alveolar and airway spaces. Additional UV-RSV control group was included. Representative histograms are included in the bottom panel. N=4-8. Each data point is an individual mouse. Each experiment was repeated at least once. Statistical significance was determined by performing One-Way ANOVA with Holm post-hoc.

### Enhanced *in vivo* responsiveness to LPS by RSV-experienced LECs

Considering the long-term epigenetic changes observed following RSV infection, we investigated if LEC responses to immunogens are altered. Lipopolysaccharide (LPS) or PBS was given intranasally (stimulation) to mice 28 days after RSV or PBS administration (conditioning), followed by transcriptome and protein analysis of LECs (**Figure 3A**). LPS has well described effects on LECs (52–54); triggering proinflammatory transcriptional programmes through TLR4/CD14/MD-2 receptor signalling, similar to RSV F protein (55), and it is a major component of house dust mite (HDM) (56), a major respiratory allergen.

**Figure 3.**
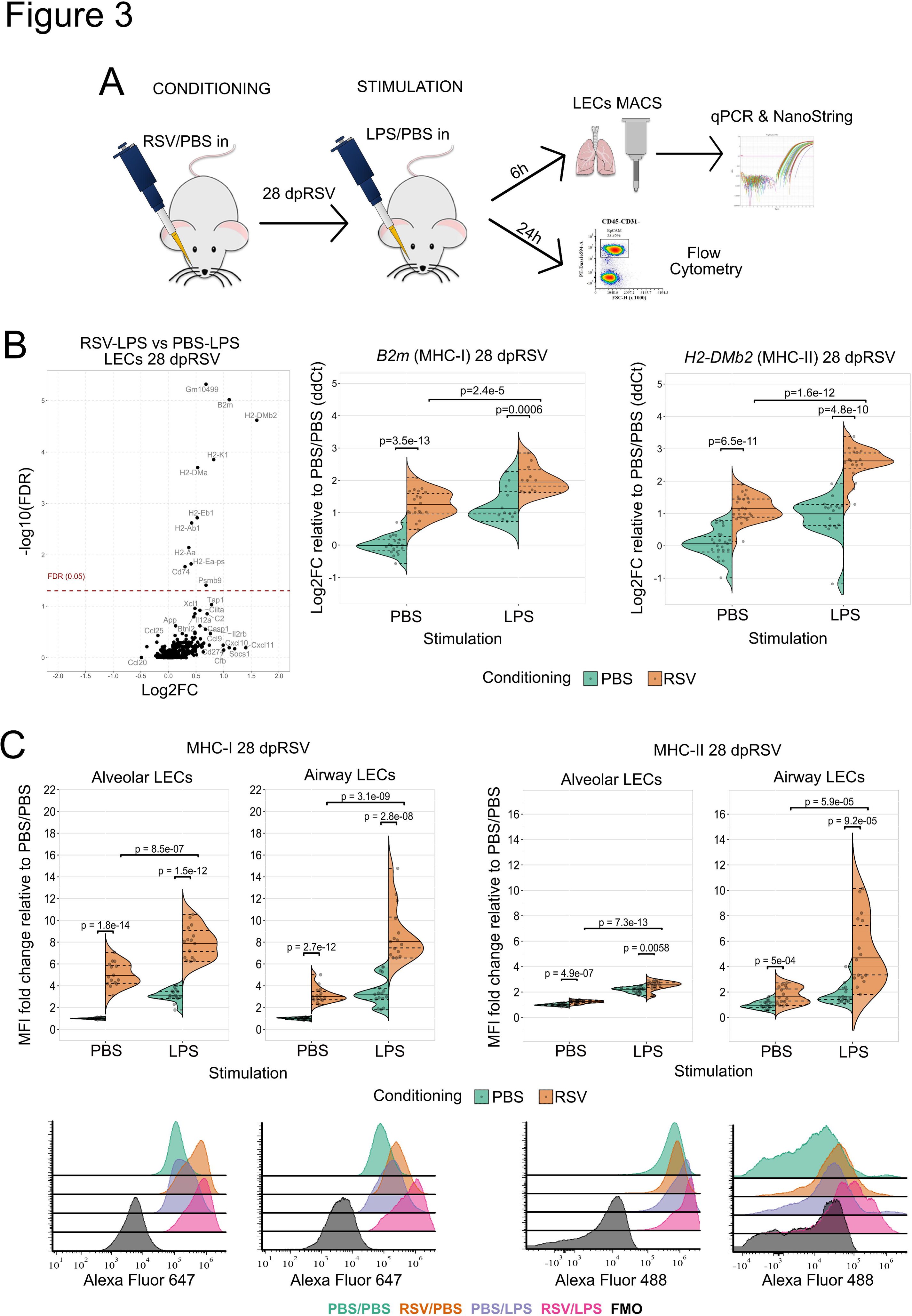
Intranasal LPS administration 28dpRSV results in an enhanced expression of MHC-I and MHC-II on LECs. (**A**) 28 days after administering RSV or PBS to 6-week-old female BALB/c mice 10ug of LPS or PBS were administered intranasally. Either 6h or 24h later mice were culled for transcriptomic or flow cytometry analysis respectively. (**B**) Left panel - volcano plot of differentially expressed LEC genes 28dpRSV identified by NanoString nCounter mouse Immunology panel in RSV-LPS groups as compared to PBS-LPS group. X axis shows a log2 fold-change in expression, while y axis shows a -log10 FDR values. NanoString n=6 per group, each sample is a pool of two mice, two independent experiments. Centre and right panels - representative MHC-I and MHC-II-related targets validated using duplex TaqMan qPCR. *Oaz1* was used as endogenous control for *H2-DMb2* while *Gapdh* was used as endogenous control for *B2m*. Treatment on day 0 (conditioning) is colour coded, PBS – green and RSV – orange. Treatment on day 28 (Stimulation with PBS or LPS) is labelled on x axis. N=12-24 per group, four independent experiments. (**C**) Relative flow cytometry quantification of MHC-I and MHC-II expression on LECs 28dpRSV stratified based on EpCAM expression which corresponds to alveolar and airway spaces. Representative histograms are included in the bottom panel. Treatment on day 0 (conditioning) is colour coded, PBS – green and RSV – orange. Treatment on day 28 (Stimulation with PBS or LPS) is labelled on x axis. N=15 per group, four independent experiments. Two-Way ANOVA with Holm post-hoc was performed to determine statistical significance.

Following 6h *in vivo* LPS or PBS stimulation, the transcriptional response of isolated LECs was assessed using NanoString (**Figure 3B**). Virtually all significantly upregulated genes (RSV-LPS vs PBS-LPS) were related to MHC biology and the increased expression of *B2m* (MHC-I) and *H2-DMb2* (MHC-II) was validated by qPCR (**Figure 3B**). Next, we used flow cytometry to assess the expression of MHC molecules on LECs 24h after LPS administration, at 28dpRSV. Considering the reports that TLR4, a LPS sensor, is upregulated on LECs following RSV infection (57) we assessed its expression using flow cytometry 28dpRSV, to ensure there are no differences compared to control (**Figure S8**). Corroborating the qPCR data, RSV conditioning enhanced the response to LPS, with increased MHC-I and MHC-II expression in both airway LECs and alveolar LECs (**Figure 3C**).

### LECs show augmented antigen uptake, processing and presentation 28dpRSV

Across CUT&RUN and NanoString data sets, we identified several targets associated with antigen processing and trafficking at 28dpRSV, namely *Psmb9*, *Lap3*, *Tap1* and *Xcl1.* The upregulation of *Psmb9 (*20S immunoproteasome subunit) and *Tap1* (transporter molecule responsible for trafficking processed antigens (58)) was further validated by qPCR 28dpRSV and was not affected by LPS stimulation (**Figure 4A**). This led us to hypothesise that either antigen uptake rate and/or antigen processing rate in LECs may be altered 28dpRSV.

**Figure 4.**
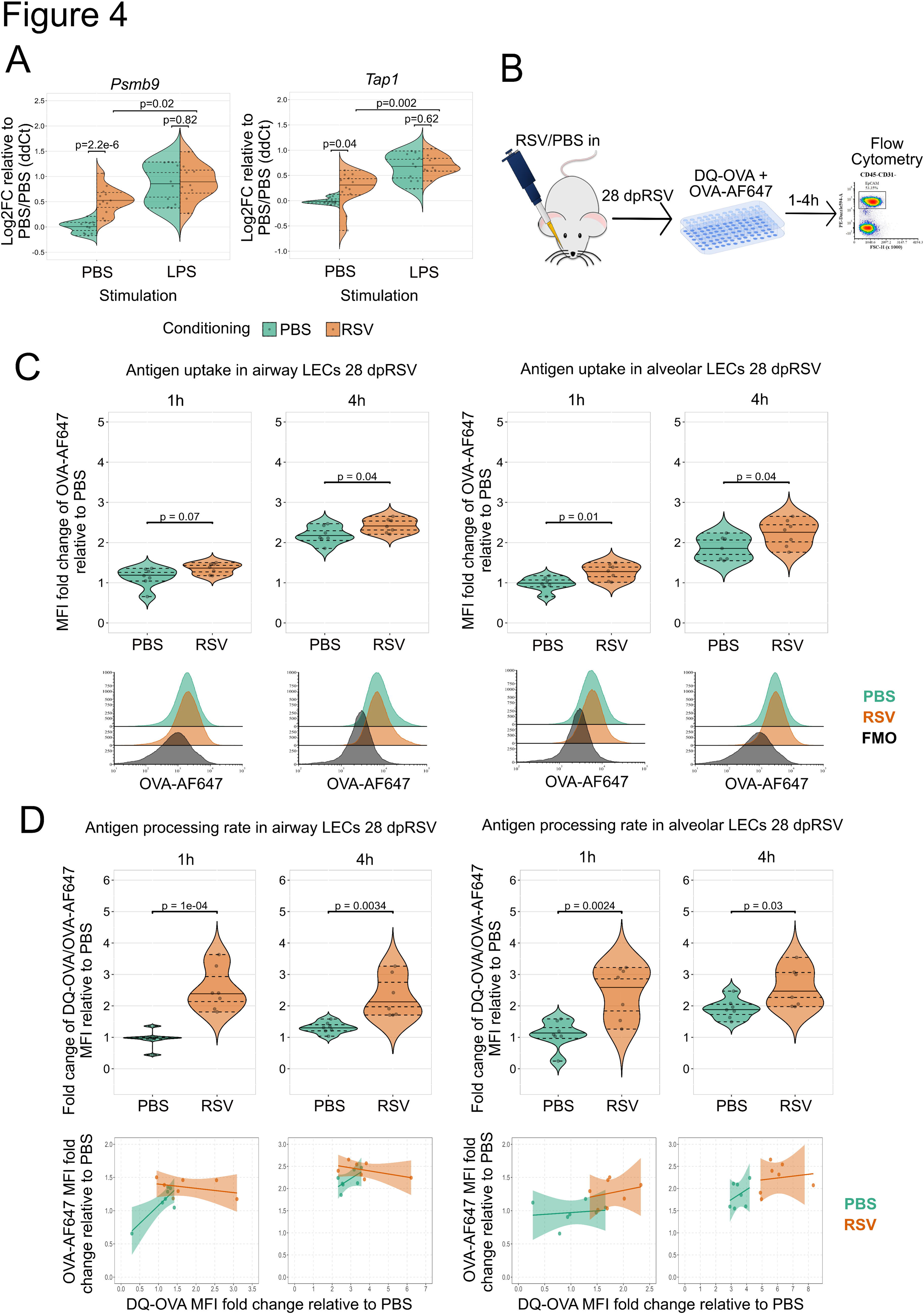
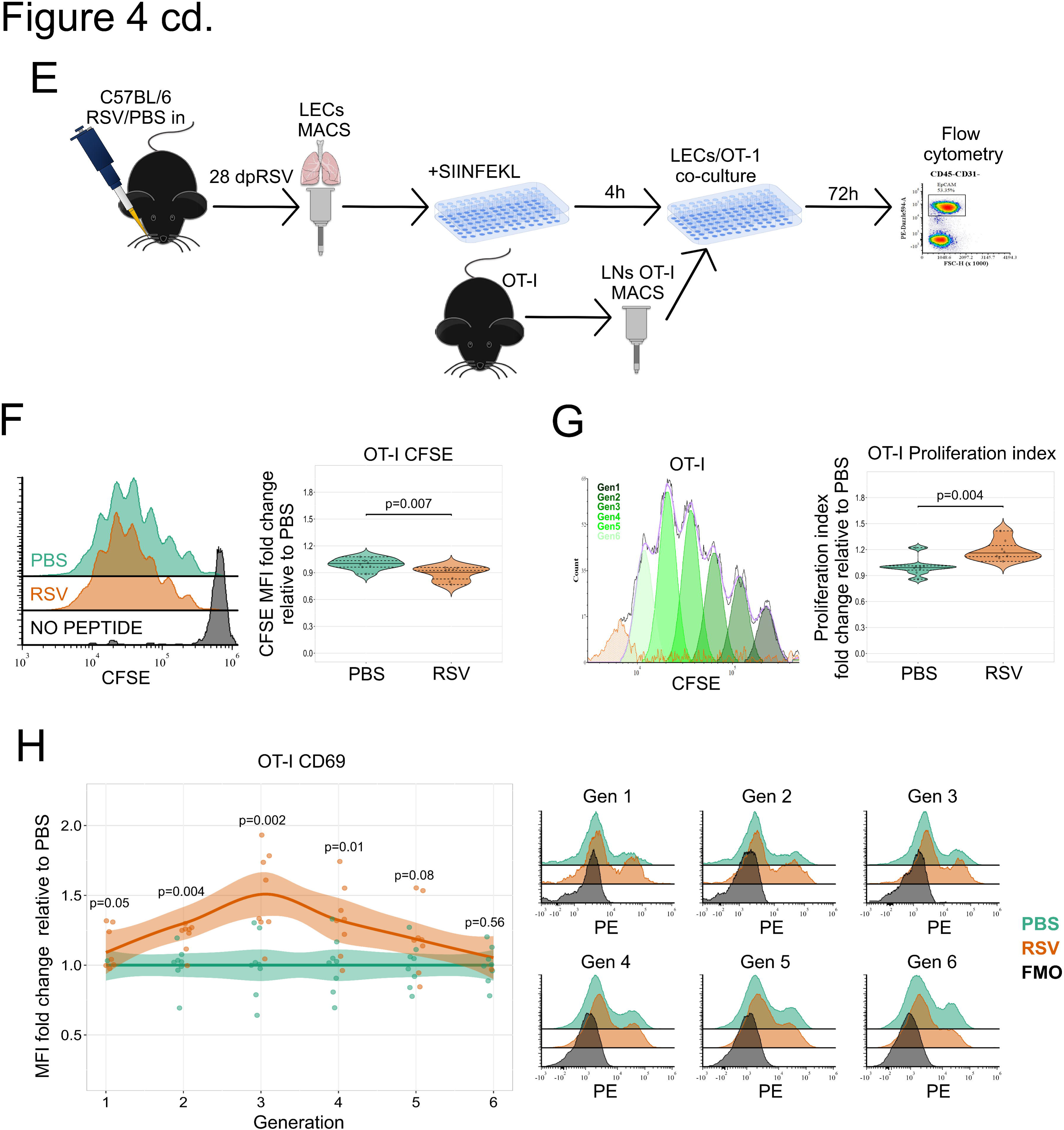
RSV infection results in enhanced LECs antigen uptake, processing and presentation. (**A**) Duplex TaqMan qPCR of *Psmb9* and *Tap1* with *Oaz1* as endogenous control in MACS-sorted LECs 28dpRSV. (**B**) Experimental design for antigen uptake and antigen processing assessment. Mice were administered RSV or PBS intranasally, followed by administration of PBS or LPS 28 days later. Mice were culled and lung single cell suspension was placed in serum-free media with DQ-OVA and OVA-AF647. After 1h or 4h cells were collected and stained for flow cytometry analysis. (**C**) Relative flow cytometry quantification of antigen uptake as determined by intracellular OVA-AF647 signal in LECs stratified based on alveolar and airway spaces. Representative histograms in the panel on the right. (**D**) Relative flow cytometry quantification of antigen processing as determined by a ratio of intracellular DQ-OVA and OVA-AF647 signal in LECs stratified based on alveolar and airway spaces. OVA-AF647/DQ-OVA bivariate plots in the panel at the bottom (**E**) Experimental design for assessment of antigen presentation. C57BL/6 mice were administered RSV or PBS intranasally. 28 days later mice were culled and LECS were MACS-sorted. Sorted LECs were placed in an in vitro culture with SIINFEKL OVA peptide and with/without LPS for 4h. In the meantime, lymph nodes were harvested from OT-I mice and OT-I CD8 T-cells were MACS sorted. OT-I cells were labelled with proliferation dye CFSE. LECs and OT-I cells were then co-cultured for 72h, followed by flow cytometry analysis. (**F**) Relative flow cytometry analysis of OT-I proliferation using CFSE dye. Left panel – representative histograms. Right panel – relative quantification of CFSE decrease. N=8 per experimental group, two independent experiments. (**G**) Relative flow cytometry analysis of OT-I proliferation using FCS express 7 proliferation index. Left panel – automated identification of OT-I generations. Right panel - relative OT-I proliferation index. N=8 per experimental group, two independent experiments. (**H**) Relative flow cytometry analysis of CD69 MFI in OT-I across six generations 72h after co-culture start with SIINFEKL-fed LECs. N=8 per experimental group, two independent experiments. PBS – green and RSV – orange. One-Way ANOVA with Holm post-hoc was performed to determine statistical significance.

To test this hypothesis, we infected or mock infected mice, followed by isolation and culture of LECs 28dpRSV in the presence of fluorescent ovalbumin (OVA)-AF647, used to indicate antigen uptake, and BODIPY FL dye-containing DQ-OVA (**Figure 4B**) which fluoresces proportionally to its proteolytic cleavage (59). Thus, the ratio of DQ-OVA and OVA-AF647 reflects the rate of antigen processing while controlling for potential differences in antigen uptake. First, we assessed the ability of LECs to take up OVA-AF647 antigen over 1h or 4h with data stratified to alveolar and airway LECS (**Figure 4C**). Increased antigen uptake was observed after RSV conditioning compared to PBS controls across both epithelial subsets and timepoints. Similarly, we assessed antigen processing rates in alveolar and airway LECs after 1h or 4h (**Figure 4D**). Again, we observed that the antigen processing rate is elevated 28dpRSV across both cell types.

Next, to test if long-term upregulation of MHC molecules results in enhanced antigen presentation, we measured proliferation of transgenic OVA-specific OT mouse T cells in response to OVA-peptide presented by LECs. Due to MHC restriction (60) and the C57BL/6J genetic background of OT mice (61) we needed to test if the effects of RSV infection are recapitulated in C57BL/6J LECs. Similar to BALB/c mice, the expression of MHC-I in C57BL/6 mice remained highly upregulated 28dpRSV, however, no long-term expression of MHC-II was observed **(Figure S9**).

28 days after the inoculation of C57BL/6J mice with RSV, we harvested LECs and cultured them with or without OVA peptide recognised by OT-I CD8 T-cells (SIINFEKL) and LPS. After 4h LECs were washed and put in a co-culture with OT-I cells labelled with CFSE proliferation dye (Figure 4E). After 72h, OT-I cells proliferated more when exposed to LECs that previously experienced RSV as indicated by a decrease in CFSE signal (**Figure 4F**) and an increase in proliferation index (**Figure 4G**). Additionally, CD69, a T-cell activation marker, mirrored the LEC MHC expression pattern in the initial generations, with significant upregulation in the RSV group (**Figure 4H**).

## Discussion

Here, we demonstrate that RSV infection results in long-term epigenetic changes in LECs, including in regulatory elements of multiple genes related to MHC molecules and antigen processing. Furthermore, we demonstrate that 28dpRSV LECs maintain high expression of MHC-I and MHC-II at transcriptional and protein levels. The expression of both MHC classes in LECs can be further enhanced upon subsequent exposure to LPS. Observed epigenetic, transcriptomic and protein changes seem to be functional, as we observe that 28dpRSV LECs maintain elevated antigen uptake and processing following incubation with OVA-AF647 and DQ-OVA. Importantly, these changes are associated with an increased propensity for T-cells to proliferate and become activated when antigen is presented by LECs that experienced RSV infection 28 days earlier.

Our findings have a twofold consequence. On one hand increased expression of MHC-I following RSV may enhance protection against subsequent intracellular pathogens. Conversely, the enhanced expression of MHC-II may lead to increased presentation of microbial antigens or allergens to CD4 T-cells. Indeed, mounting evidence suggests that the severity of influenza infection is reduced if mice were infected with RSV 3-4 weeks prior as observed by reduced morbidity, mortality, lung pathology and immune cell infiltration (8,61). This phenomenon was shown to be independent of antibody-mediated protection or cross-protecting lung-resident memory T-cells (T_RM_) (62). Here, we suggest that the observed protection could be a result of long-term enhancement of endogenous antigen presentation by LECs.

The importance of MHC antigen presentation during viral infection is underscored by the observation that many viruses including SARS-CoV-2, influenza virus, cytomegalovirus, Epstein-Barr virus, HIV, rotavirus or hepatitis B virus, actively interfere with MHC-I expression, antigen loading and presentation, including in epithelial cells (63–69). RSV infection, on the other hand, as shown here *in vivo* in mice, as well as by other researchers in human lung epithelial cell lines (70,71) upregulates the expression of MHC. Interestingly, rhinovirus, another virus which is associated with allergic airway disease development and exacerbation (72,73), also upregulates MHC in respiratory epithelial cells (74).

While enhanced LEC antigen presentation may contribute to allergic asthma exacerbation; it is unlikely that heightened LEC allergen presentation alone will lead to the development of allergic airway disease. Rather, other confounding factors like genetic predisposition (MHC alleles) or environmental factors all contribute to development of allergic airway disease. HLA genes are the most polymorphic genes found in humans (75), with more than 35,000 alleles identified as of 2023 (76). It is therefore likely that only specific combinations of HLA alleles may increase the risk for asthma development, particularly considering that different MHC alleles have varying affinity for different peptides (77). Indeed, it has been shown that different class II HLA genes have an impact on the course of asthma development (78). Meanwhile, several independent genome-wide association studies (GWAS) identified HLA-DQ as an allele predisposing to asthma development (75,79,80). Nevertheless, in light of a report that lymph nodes are not essential for clonal expansion of allergen-specific CD4 T-cells in the murine lung (81), it is conceivable that long-term MHC upregulation on LECs alone might suffice for the expansion of allergen-specific lung-resident T-cell populations, and thus through repeated allergen exposure contribute to development of allergic airway disease.

Considering the increase in MHC levels (e.g. following RSV infection) resulting in enhanced antigen presentation, it is possible that the presence of environmental LPS may contribute to MHC-mediated enhanced allergen presentation and thus allergic airway disease exacerbation in susceptible individuals. Multiple studies utilizing various experimental systems demonstrated that LPS exposure results in allergic airway exacerbation (82–84), while another study also demonstrated that repeated LPS administration 35-41 days after RSV infection results in aggravated inflammatory response and airway hyper-responsiveness (85).

While MHC-I and MHC-II upregulation is generally IFNγ-mediated (86–88), MHC-I upregulation following RSV has been demonstrated to be primarily type I interferon (IFN-I) mediated (71,89). Nevertheless, it is unlikely that IFNs alone are responsible for the observed phenotype. While C57BL/6 mice prototypically exhibit type 1 innate immune responses (T1), BALB/c mice tend to exhibit a type 2 innate immune response (T2) (90). We observed that RSV infection upregulates MHC-II on LECs in BALB/c mice but not in C57BL/6 mice, suggesting that while general antiviral responses may be responsible for upregulation of MHC-I, a T2 cytokine response may be responsible for upregulation of MHC-II. This notion is further reinforced considering that T2 cytokines like IL-13 or IL-33 correlate with RSV disease severity (45,91), which in our model correlates with long-term expression levels of MHC. Similarly, several T2 cytokines including IL-2, IL-4 or IL-13 induce MHC-II expression (92,93).

In summary, using a mouse model we have shown that RSV infection results in long-term epigenetic and transcriptomic changes to lung epithelial cells which result in increased antigen presentation. Our findings increase understanding of epithelial cell memory after infection and could explain severity modulation of subsequent LRTI as well as the association between LRTI and the development of allergic airway disease.

## Author Contributions

Conceptualization: PPJ, HJM, WTJ, EWR and JS. Investigation: PPJ, WTJ and MOB. Formal analysis: PPJ, WTJ, MV. Writing – original draft: PPJ. Writing – review and editing: PPJ, WTJ, MOB, CC, MV, CL, CB, RI, EWR, HJM and JS. Funding acquisition: JS. Project administration: JS. EWR provided the OT-I mice.

