## Supplementary figures for "Epithelial memory after respiratory viral infection results in long-lasting enhancement of antigen presentation"

### Figure S1

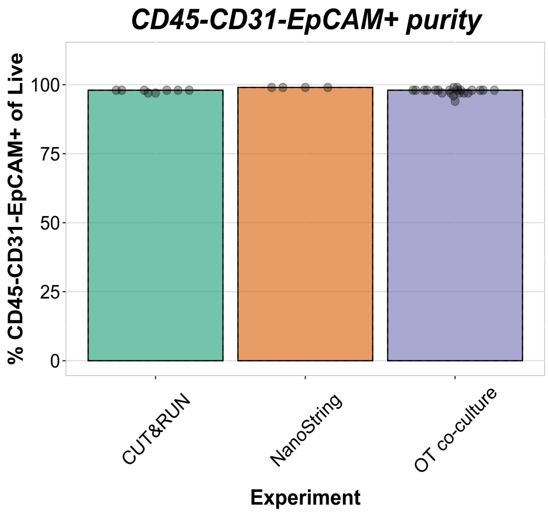


### Figure S2

PBS H3K4me3

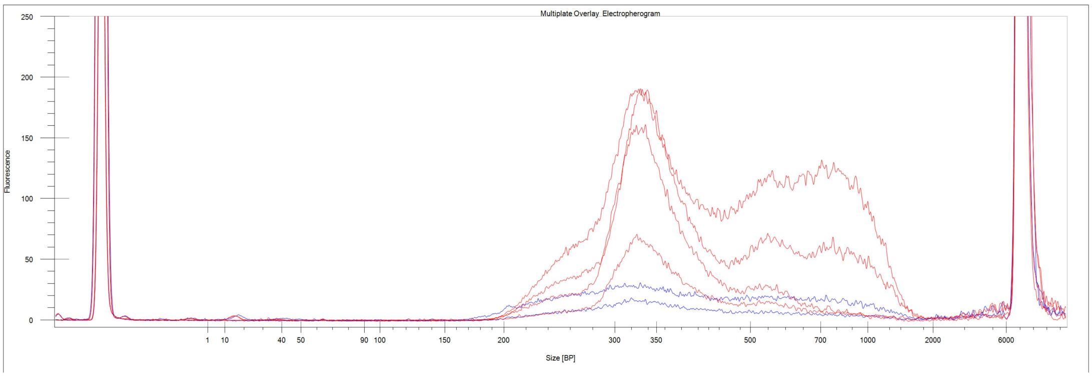


RSV H3K4me3

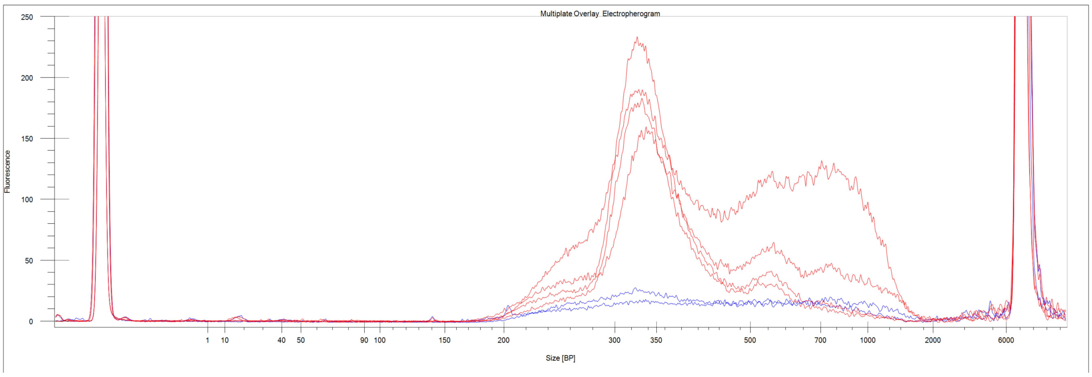


PBS H3K27ac

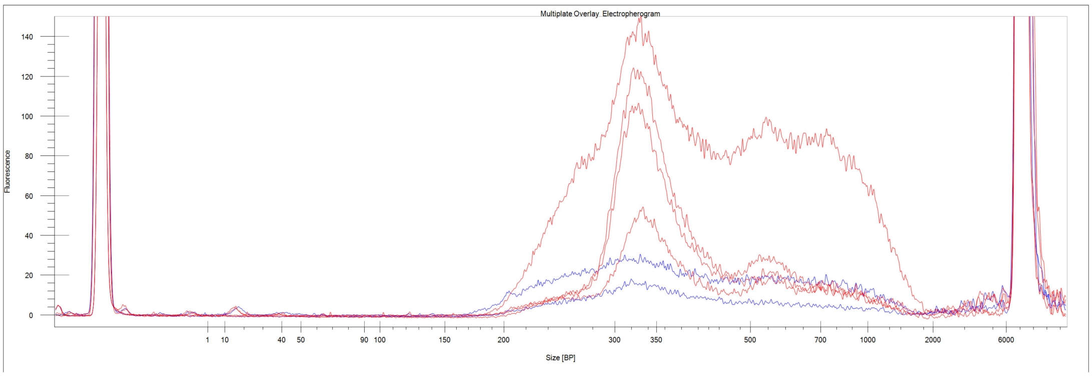


RSV H3K4me3

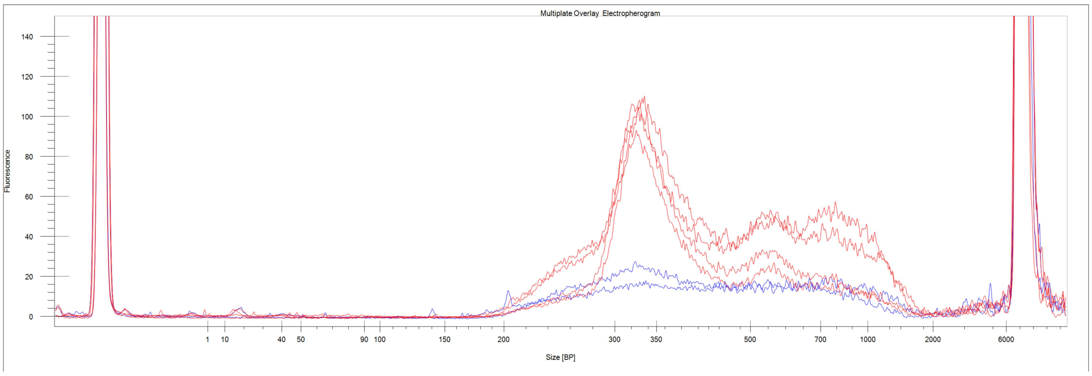

### Figure S3

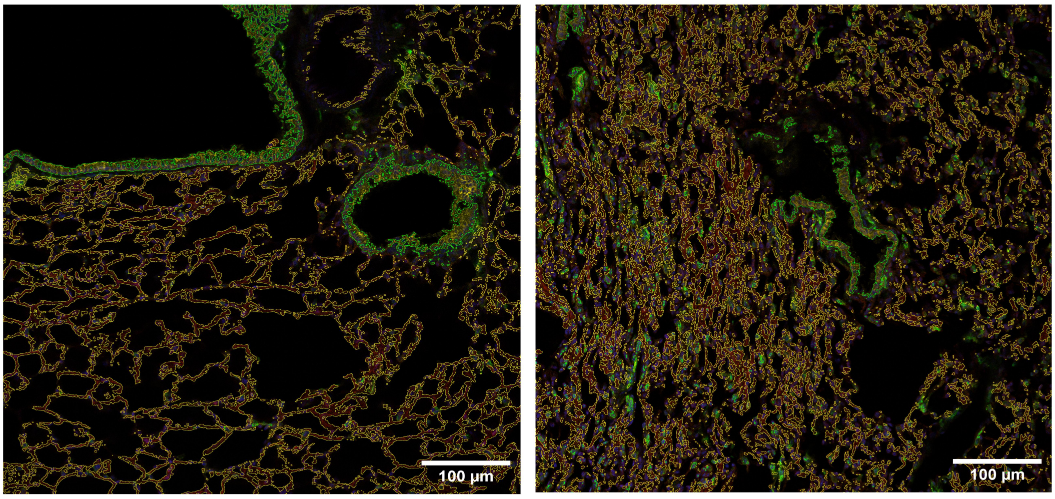


### Figure S4

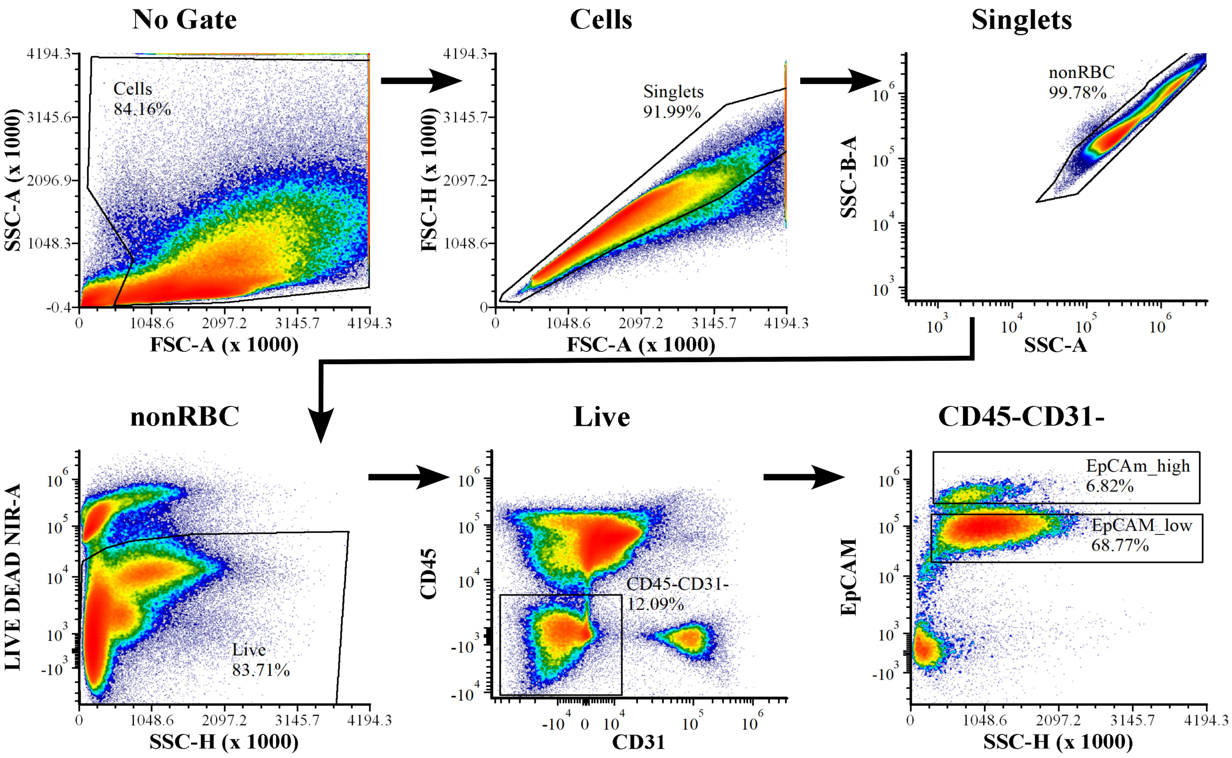

### Figure S5

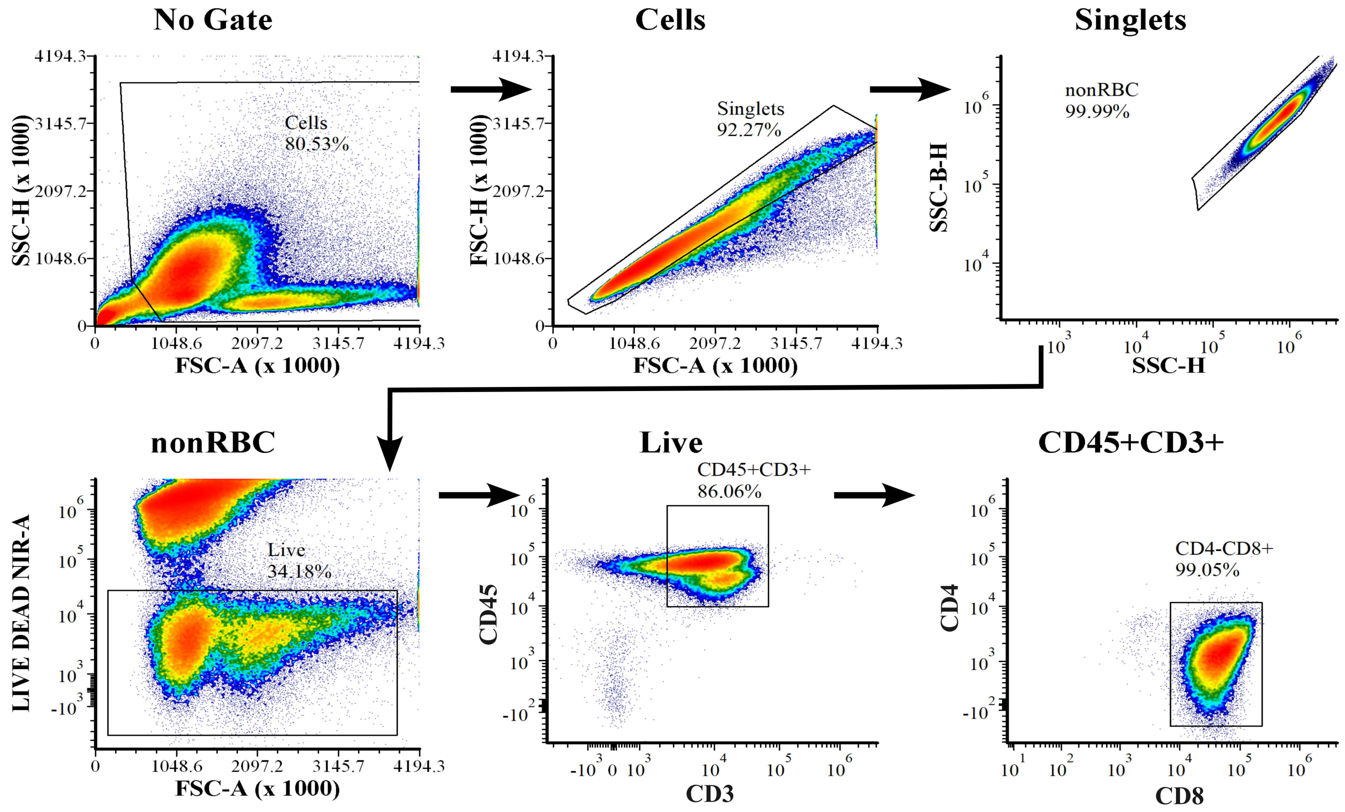


### Figure S6

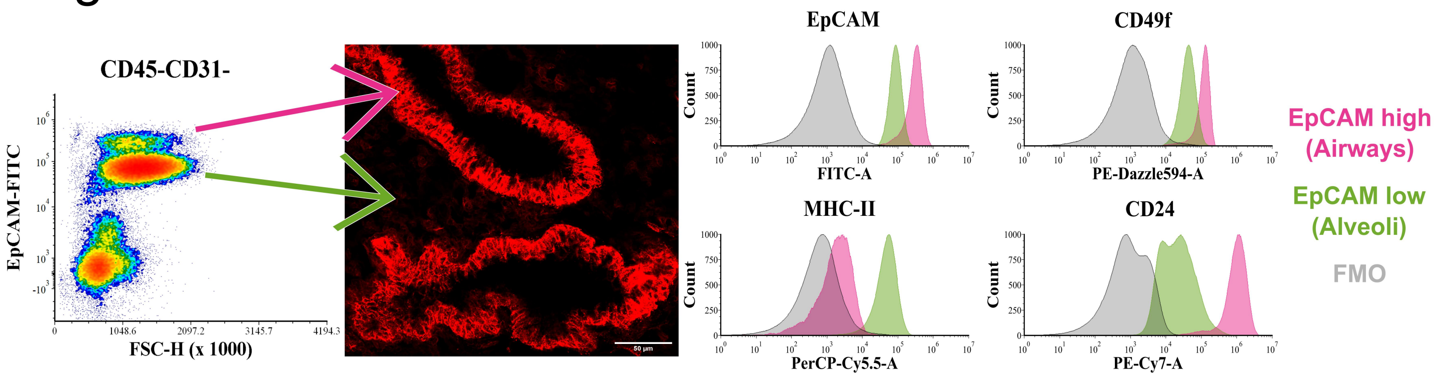

### Figure S7

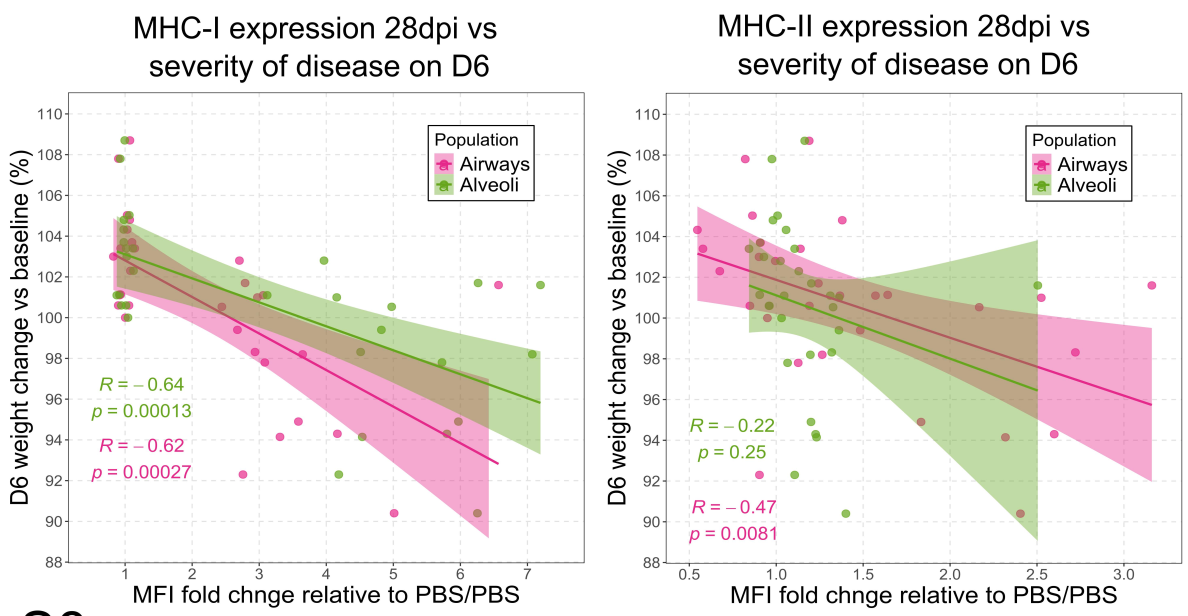


### Figure S8

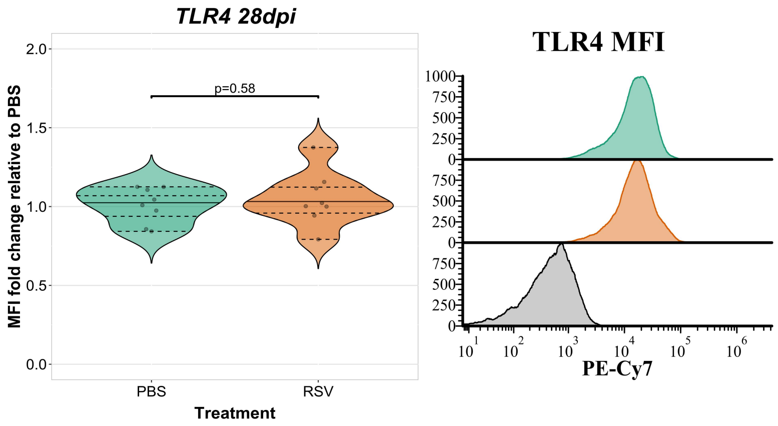


### Figure S9

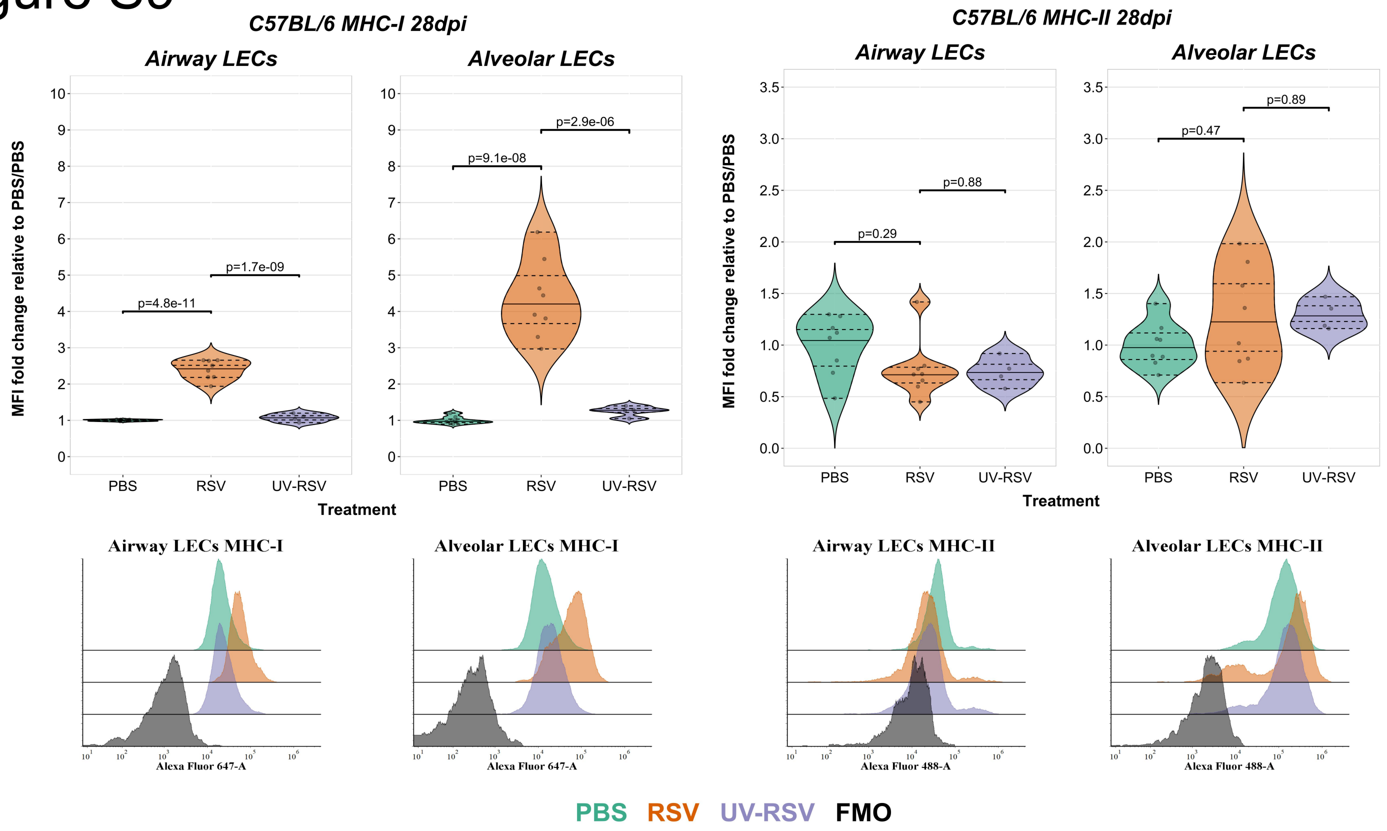

### Table S1

| Gene | GroupDif | GroupSD | **Stability score** |
|---|---|---|---|
| *Oaz1* | 0.19 | 0.33 | **0.13** |
| *Gapdh* | 0.43 | 0.58 | **0.21** |
| *Rpl37a* | 0.71 | 0.65 | **0.24** |
| *18S* | 1.09 | 0.86 | **0.28** |

### Table S2

| Histone modification | SYMBOL | Conc_RSV | Conc_PBS | Fold | p.value | FDR | Genomic annotation | gene ID | ENSEMBL |
|---|---|---|---|---|---|---|---|---|---|
| H3K4me3 | Iigp1 | 11.09 | 9.69 | 1.377542136 | 1.60E-30 | 2.06E-26 | Promoter (<=1kb) | 60440 | ENSMUSG00000054072 |
| H3K4me3 | Oas1a | 9.69 | 7.99 | 1.65259025 | 8.10E-28 | 5.20E-24 | Promoter (<=1kb) | 246730 | ENSMUSG00000052776 |
| H3K4me3 | Oas1g | 9.60 | 8.59 | 0.998866862 | 8.93E-20 | 3.82E-16 | Promoter (<=1kb) | 23960 | ENSMUSG00000066861 |
| H3K4me3 | Ifit1bl2 | 10.43 | 9.88 | 0.519410604 | 6.19E-12 | 1.99E-08 | Promoter (<=1kb) | 112419 | ENSMUSG00000067297 |
| H3K4me3 | Gm5431 | 10.17 | 9.04 | 1.08958445 | 1.87E-11 | 4.79E-08 | Promoter (<=1kb) | 432555 | ENSMUSG00000058163 |
| H3K4me3 | Ifi47 | 11.99 | 11.55 | 0.411510731 | 8.88E-11 | 1.90E-07 | Promoter (<=1kb) | 15953 | ENSMUSG00000078920 |
| H3K4me3 | Gbp9 | 9.72 | 8.66 | 1.016247832 | 9.27E-09 | 1.70E-05 | Promoter (<=1kb) | 236573 | ENSMUSG00000029298 |
| H3K4me3 | H2-K2 | 10.82 | 10.46 | 0.308805978 | 5.14E-08 | 7.82E-05 | Promoter (<=1kb) | 630499 | ENSMUSG00000121503 |
| H3K4me3 | 2700069I18Rik | 10.23 | 10.58 | -0.285249807 | 5.49E-08 | 7.82E-05 | Promoter (<=1kb) | 72608 | NA |
| H3K4me3 | 9330175E14Rik | 9.88 | 9.12 | 0.687577858 | 1.43E-07 | 0.000183545 | Promoter (<=1kb) | 320377 | NA |
| H3K4me3 | Sorbs1 | 9.79 | 9.39 | 0.330893614 | 4.20E-07 | 0.00048974 | Intron | 20411 | ENSMUSG00000025006 |
| H3K4me3 | Dhx58 | 9.77 | 9.22 | 0.458356247 | 5.62E-07 | 0.00060041 | Promoter (<=1kb) | 80861 | ENSMUSG00000017830 |
| H3K4me3 | Spats2l | 11.00 | 10.71 | 0.205803055 | 2.14E-06 | 0.00201797 | Promoter (<=1kb) | 67198 | ENSMUSG00000038305 |
| H3K4me3 | Frzb | 10.54 | 10.76 | -0.163336089 | 2.34E-06 | 0.00201797 | Promoter (<=1kb) | 20378 | ENSMUSG00000027004 |
| H3K4me3 | H2-D1 | 12.22 | 11.94 | 0.201098376 | 2.36E-06 | 0.00201797 | Promoter (<=1kb) | 14964 | ENSMUSG00000073411 |
| H3K4me3 | H2-T22 | 11.39 | 10.87 | 0.385244993 | 3.01E-06 | 0.002409672 | Promoter (<=1kb) | 15039 | ENSMUSG00000056116 |
| H3K4me3 | Tuba8 | 10.72 | 10.47 | 0.172048398 | 5.75E-06 | 0.004338839 | Promoter (<=1kb) | 53857 | ENSMUSG00000030137 |
| H3K4me3 | Jarid2 | 10.11 | 10.42 | -0.199222683 | 1.07E-05 | 0.007640145 | Promoter (<=1kb) | 16468 | ENSMUSG00000038518 |
| H3K4me3 | Ptprd | 10.72 | 10.97 | -0.165996979 | 1.24E-05 | 0.007988958 | Promoter (<=1kb) | 19266 | ENSMUSG00000028399 |
| H3K4me3 | Ciart | 10.31 | 9.89 | 0.258762407 | 1.25E-05 | 0.007988958 | Promoter (<=1kb) | 229599 | ENSMUSG00000038550 |
| H3K4me3 | Tab2 | 11.44 | 11.61 | -0.128948083 | 1.46E-05 | 0.008938019 | Promoter (<=1kb) | 68652 | ENSMUSG00000015755 |
| H3K4me3 | Ptprq | 9.76 | 10.08 | -0.183194781 | 2.86E-05 | 0.016649103 | Distal Intergenic | 237523 | ENSMUSG00000035916 |
| H3K4me3 | Rcan1 | 11.03 | 10.82 | 0.140214634 | 3.46E-05 | 0.019317788 | Promoter (<=1kb) | 54720 | ENSMUSG00000022951 |
| H3K4me3 | B2m | 11.57 | 11.20 | 0.188822989 | 4.64E-05 | 0.024813953 | Promoter (<=1kb) | 12010 | ENSMUSG00000060802 |
| H3K4me3 | Cald1 | 10.46 | 10.19 | 0.148853144 | 5.56E-05 | 0.028518044 | Exon | 109624 | ENSMUSG00000029761 |
| H3K4me3 | Pcdh9 | 11.13 | 11.38 | -0.144280672 | 6.58E-05 | 0.03247394 | Promoter (<=1kb) | 211712 | ENSMUSG00000055421 |
| H3K4me3 | Oasl2 | 10.32 | 9.76 | 0.196252947 | 7.37E-05 | 0.033821296 | Promoter (<=1kb) | 23962 | ENSMUSG00000029561 |
| H3K4me3 | AW112010 | 9.94 | 9.35 | 0.199918703 | 7.45E-05 | 0.033821296 | Promoter (<=1kb) | 107350 | ENSMUSG00000075010 |
| H3K4me3 | H2-Ab1 | 10.96 | 10.55 | 0.172218535 | 7.65E-05 | 0.033821296 | Promoter (<=1kb) | 14961 | ENSMUSG00000073421 |
| H3K4me3 | Ifi44 | 9.86 | 9.46 | 0.166103544 | 8.43E-05 | 0.036055844 | Promoter (<=1kb) | 99899 | ENSMUSG00000028037 |
| H3K4me3 | Rexo1 | 11.20 | 11.00 | 0.125997305 | 9.62E-05 | 0.039794855 | Promoter (<=1kb) | 66932 | ENSMUSG00000047417 |
| H3K27ac | Iigp1 | 11.24 | 9.62 | 1.562787554 | 8.42E-29 | 1.48E-24 | Promoter (<=1kb) | 60440 | ENSMUSG00000054072 |
| H3K27ac | Ddx60 | 9.33 | 7.84 | 1.407686453 | 1.80E-24 | 1.58E-20 | Promoter (<=1kb) | 234311 | ENSMUSG00000037921 |
| H3K27ac | Gbp9 | 9.48 | 8.01 | 1.370888755 | 8.91E-17 | 5.23E-13 | Promoter (<=1kb) | 236573 | ENSMUSG00000029298 |
| H3K27ac | Igtp | 9.16 | 7.57 | 1.449220429 | 1.42E-14 | 6.23E-11 | Promoter (<=1kb) | 16145 | ENSMUSG00000078853 |
| H3K27ac | B2m | 10.13 | 9.02 | 1.038191433 | 4.22E-13 | 1.49E-09 | Promoter (<=1kb) | 12010 | ENSMUSG00000060802 |
| H3K27ac | Gbp8 | 8.52 | 6.85 | 1.481282945 | 2.70E-11 | 7.93E-08 | Promoter (<=1kb) | 76074 | ENSMUSG00000034438 |
| H3K27ac | Stat1 | 8.90 | 7.91 | 0.910862368 | 4.77E-11 | 1.20E-07 | Promoter (<=1kb) | 20846 | ENSMUSG00000026104 |
| H3K27ac | H2-Eb1 | 8.32 | 6.72 | 1.452174413 | 2.21E-10 | 4.87E-07 | Promoter (<=1kb) | 14969 | ENSMUSG00000060586 |
| H3K27ac | Prpf40a | 9.56 | 8.50 | 0.952887182 | 1.23E-09 | 2.16E-06 | Promoter (<=1kb) | 56194 | ENSMUSG00000061136 |
| H3K27ac | Ifi47 | 12.12 | 11.80 | 0.271844774 | 1.38E-09 | 2.20E-06 | Promoter (<=1kb) | 15953 | ENSMUSG00000078920 |
| H3K27ac | Rp9 | 8.52 | 7.66 | 0.755081135 | 4.84E-09 | 7.10E-06 | Promoter (<=1kb) | 55934 | ENSMUSG00000032239 |
| H3K27ac | Clcn3 | 10.03 | 9.54 | 0.393629029 | 8.43E-09 | 1.14E-05 | Promoter (<=1kb) | 12725 | ENSMUSG00000004319 |
| H3K27ac | H2-Ab1 | 8.59 | 7.07 | 1.312938172 | 9.08E-09 | 1.14E-05 | Promoter (<=1kb) | 14961 | ENSMUSG00000073421 |
| H3K27ac | Tbl1xr1 | 10.59 | 10.93 | -0.275526686 | 4.15E-08 | 4.87E-05 | 5' UTR | 81004 | ENSMUSG00000027630 |
| H3K27ac | Spats2l | 10.98 | 10.51 | 0.370146046 | 7.49E-08 | 8.25E-05 | Promoter (<=1kb) | 67198 | ENSMUSG00000038305 |
| H3K27ac | Ccng2 | 9.97 | 9.36 | 0.474725443 | 8.13E-08 | 8.42E-05 | Promoter (<=1kb) | 12452 | ENSMUSG00000029385 |
| H3K27ac | Brwd3 | 9.22 | 8.37 | 0.674553896 | 3.24E-07 | 0.000302857 | Promoter (<=1kb) | 382236 | ENSMUSG00000063663 |
| H3K27ac | AW112010 | 10.57 | 10.17 | 0.314035251 | 3.29E-07 | 0.000302857 | Promoter (<=1kb) | 107350 | ENSMUSG00000075010 |
| H3K27ac | H2-D1 | 9.65 | 8.54 | 0.918208757 | 3.56E-07 | 0.000302857 | Promoter (<=1kb) | 14964 | ENSMUSG00000073411 |
| H3K27ac | Zbp1 | 8.99 | 8.14 | 0.69132064 | 3.61E-07 | 0.000302857 | Promoter (<=1kb) | 58203 | ENSMUSG00000027514 |
| H3K27ac | Fam151b | 8.31 | 9.06 | -0.564270295 | 9.01E-07 | 0.000721485 | 3' UTR | 73942 | ENSMUSG00000034334 |
| H3K27ac | Gm5431 | 10.02 | 9.54 | 0.353095569 | 9.74E-07 | 0.000737092 | Promoter (<=1kb) | 432555 | ENSMUSG00000058163 |
| H3K27ac | Mia | 9.56 | 8.96 | 0.430899038 | 1.00E-06 | 0.000737092 | Promoter (<=1kb) | 12587 | ENSMUSG00000089661 |
| H3K27ac | Zxdc | 9.20 | 9.61 | -0.294318248 | 1.86E-06 | 0.001274843 | 3' UTR | 80292 | ENSMUSG00000034430 |
| H3K27ac | Carf | 8.44 | 7.53 | 0.658420834 | 1.88E-06 | 0.001274843 | Promoter (<=1kb) | 241066 | ENSMUSG00000026017 |
| H3K27ac | Hdgf | 9.97 | 8.95 | 0.75499166 | 2.46E-06 | 0.001601168 | Promoter (<=1kb) | 15191 | ENSMUSG00000004897 |
| H3K27ac | Spag9 | 10.26 | 9.76 | 0.344176796 | 2.62E-06 | 0.00165044 | Promoter (<=1kb) | 70834 | ENSMUSG00000020859 |
| H3K27ac | Cap1 | 10.43 | 9.89 | 0.362499209 | 3.33E-06 | 0.002022642 | Promoter (<=1kb) | 12331 | ENSMUSG00000028656 |
| H3K27ac | Caprin1 | 9.86 | 9.26 | 0.396977602 | 3.64E-06 | 0.002137967 | Promoter (<=1kb) | 53872 | ENSMUSG00000027184 |
| H3K27ac | 1700108F19Rik | 8.21 | 8.77 | -0.380037272 | 3.78E-06 | 0.002144844 | Distal Intergenic | 73272 | ENSMUSG00000101009 |
| H3K27ac | Gm12185 | 9.06 | 8.42 | 0.448034704 | 4.18E-06 | 0.002298405 | Promoter (<=1kb) | 620913 | ENSMUSG00000048852 |
| H3K27ac | Qars | 9.17 | 8.40 | 0.508643967 | 5.71E-06 | 0.002981191 | Promoter (<=1kb) | 97541 | ENSMUSG00000032604 |
| H3K27ac | Erp29 | 9.31 | 8.83 | 0.328504888 | 5.90E-06 | 0.002981191 | Promoter (<=1kb) | 67397 | ENSMUSG00000029616 |
| H3K27ac | Klhl21 | 10.54 | 10.10 | 0.30274333 | 5.93E-06 | 0.002981191 | Promoter (<=1kb) | 242785 | ENSMUSG00000073700 |
| H3K27ac | Dnajc1 | 10.02 | 10.37 | -0.255227412 | 6.41E-06 | 0.003134302 | Intron | 13418 | ENSMUSG00000026740 |
| H3K27ac | Tcta | 10.63 | 10.29 | 0.254327877 | 7.07E-06 | 0.003363751 | Promoter (<=1kb) | 102791 | ENSMUSG00000039461 |
| H3K27ac | Xrcc1 | 9.49 | 8.70 | 0.497903077 | 7.57E-06 | 0.003488983 | Promoter (<=1kb) | 22594 | ENSMUSG00000051768 |
| H3K27ac | Wbp2 | 9.76 | 9.02 | 0.464559185 | 7.73E-06 | 0.003488983 | Promoter (<=1kb) | 22378 | ENSMUSG00000034341 |

### Table S2 cd.

| Histone modification | SYMBOL | Conc_RSV | Conc_PBS | Fold | p.value | FDR | Genomic annotation | gene ID | ENSEMBL |
|---|---|---|---|---|---|---|---|---|---|
| H3K27ac | Mir6381 | 10.48 | 9.89 | 0.365865443 | 8.85E-06 | 0.003893946 | Promoter (<=1kb) | 102465200 | ENSMUSG00000098871 |
| H3K27ac | Abcc5 | 8.40 | 7.38 | 0.615806542 | 1.21E-05 | 0.005146703 | Promoter (<=1kb) | 27416 | ENSMUSG00000022822 |
| H3K27ac | Lacc1 | 8.31 | 8.82 | -0.323654275 | 1.23E-05 | 0.005146703 | Distal Intergenic | 210808 | ENSMUSG00000044350 |
| H3K27ac | Il4i1b | 9.37 | 7.96 | 0.827421003 | 1.34E-05 | 0.00548406 | Promoter (<=1kb) | 100328588 | ENSMUSG00000074141 |
| H3K27ac | Cdk17 | 9.97 | 10.42 | -0.287634382 | 1.41E-05 | 0.005588339 | Intron | 237459 | ENSMUSG00000020015 |
| H3K27ac | Lnx2 | 10.00 | 9.46 | 0.33539185 | 1.43E-05 | 0.005588339 | Promoter (<=1kb) | 140887 | ENSMUSG00000016520 |
| H3K27ac | Yeats2 | 8.60 | 9.20 | -0.369181587 | 1.57E-05 | 0.006019341 | Exon | 208146 | ENSMUSG00000041215 |
| H3K27ac | Cry2 | 11.78 | 11.31 | 0.301651847 | 1.80E-05 | 0.006731881 | Promoter (<=1kb) | 12953 | ENSMUSG00000068742 |
| H3K27ac | Angpt2 | 8.49 | 9.02 | -0.327026473 | 1.90E-05 | 0.006925112 | Intron | 11601 | ENSMUSG00000031465 |
| H3K27ac | Zc3h4 | 11.51 | 11.24 | 0.205475406 | 1.93E-05 | 0.006925112 | Promoter (<=1kb) | 330474 | ENSMUSG00000059273 |
| H3K27ac | Gga1 | 9.84 | 8.97 | 0.491624414 | 2.03E-05 | 0.007132008 | Promoter (<=1kb) | 106039 | ENSMUSG00000033128 |
| H3K27ac | Slc35f3 | 8.49 | 9.10 | -0.36201081 | 2.46E-05 | 0.008483181 | Intron | 210027 | ENSMUSG00000057060 |
| H3K27ac | Ilrun | 10.08 | 9.73 | 0.249435652 | 2.53E-05 | 0.008577637 | Promoter (<=1kb) | 224647 | ENSMUSG00000056692 |
| H3K27ac | Snora33 | 8.82 | 8.01 | 0.439199343 | 2.74E-05 | 0.009089682 | Promoter (<=1kb) | 100529074 | ENSMUSG00000070063 |
| H3K27ac | Gm10575 | 10.77 | 10.13 | 0.360330583 | 3.14E-05 | 0.010232508 | Promoter (<=1kb) | 100126795 | ENSMUSG00000073787 |
| H3K27ac | Herc6 | 10.00 | 9.51 | 0.298955373 | 3.48E-05 | 0.011132232 | Promoter (<=1kb) | 67138 | ENSMUSG00000029798 |
| H3K27ac | Wfdc1 | 8.25 | 7.02 | 0.605547974 | 3.73E-05 | 0.011733442 | Exon | 67866 | ENSMUSG00000023336 |
| H3K27ac | Myo1c | 10.39 | 9.66 | 0.385957503 | 3.97E-05 | 0.012258683 | Promoter (<=1kb) | 17913 | ENSMUSG00000017774 |
| H3K27ac | Glrx2 | 8.96 | 8.42 | 0.301778864 | 4.18E-05 | 0.012677013 | Promoter (<=1kb) | 69367 | ENSMUSG00000018196 |
| H3K27ac | Acd | 8.23 | 7.46 | 0.397922322 | 4.44E-05 | 0.013049677 | Promoter (<=1kb) | 497652 | ENSMUSG00000038000 |
| H3K27ac | Mir92b | 10.11 | 9.61 | 0.295272337 | 4.47E-05 | 0.013049677 | Promoter (<=1kb) | 100124470 | ENSMUSG00000076255 |
| H3K27ac | Paxx | 10.19 | 9.76 | 0.268453517 | 4.52E-05 | 0.013049677 | Promoter (<=1kb) | 227622 | ENSMUSG00000047617 |
| H3K27ac | Bst2 | 9.11 | 8.16 | 0.471808744 | 4.66E-05 | 0.013248145 | Promoter (<=1kb) | 69550 | ENSMUSG00000046718 |
| H3K27ac | Zfp963 | 9.33 | 8.62 | 0.371133169 | 4.79E-05 | 0.013391464 | Promoter (<=1kb) | 620419 | ENSMUSG00000092260 |
| H3K27ac | Cct7 | 10.06 | 9.32 | 0.379180105 | 5.06E-05 | 0.013909359 | Promoter (<=1kb) | 12468 | ENSMUSG00000030007 |
| H3K27ac | Ankrd28 | 8.65 | 7.99 | 0.339192408 | 5.99E-05 | 0.01623362 | Promoter (<=1kb) | 105522 | ENSMUSG00000014496 |
| H3K27ac | Pcgf5 | 8.39 | 6.95 | 0.561697894 | 6.10E-05 | 0.016261777 | Promoter (1-2kb) | 76073 | ENSMUSG00000024805 |
| H3K27ac | Smarca5 | 9.80 | 9.06 | 0.364808958 | 6.26E-05 | 0.016457431 | Promoter (<=1kb) | 93762 | ENSMUSG00000031715 |
| H3K27ac | Lcorl | 8.90 | 7.51 | 0.550675117 | 6.36E-05 | 0.016459782 | Promoter (<=1kb) | 209707 | ENSMUSG00000015882 |
| H3K27ac | Pvt1 | 9.22 | 9.70 | -0.274800678 | 6.71E-05 | 0.017123695 | Intron | 19296 | ENSMUSG00000097039 |
| H3K27ac | Pigr | 8.45 | 7.66 | 0.366913739 | 7.01E-05 | 0.017289083 | Promoter (<=1kb) | 18703 | ENSMUSG00000026417 |
| H3K27ac | Otub1 | 8.30 | 7.41 | 0.414440761 | 7.12E-05 | 0.017289083 | Promoter (<=1kb) | 107260 | ENSMUSG00000024767 |
| H3K27ac | Robo1 | 7.84 | 8.46 | -0.320224971 | 7.15E-05 | 0.017289083 | Intron | 19876 | ENSMUSG00000022883 |
| H3K27ac | Adss | 10.55 | 10.23 | 0.225757438 | 7.24E-05 | 0.017289083 | Promoter (<=1kb) | 11566 | ENSMUSG00000015961 |
| H3K27ac | Slc25a17 | 11.50 | 11.75 | -0.188256378 | 7.27E-05 | 0.017289083 | Promoter (<=1kb) | 20524 | ENSMUSG00000022404 |
| H3K27ac | Dnaaf10 | 8.89 | 8.09 | 0.373186544 | 7.96E-05 | 0.018681239 | Promoter (<=1kb) | 103784 | ENSMUSG00000078970 |
| H3K27ac | Ppp2r2b | 9.73 | 10.11 | -0.240155285 | 8.41E-05 | 0.019165999 | Intron | 72930 | ENSMUSG00000024500 |
| H3K27ac | Nop16 | 9.05 | 8.40 | 0.318264948 | 8.49E-05 | 0.019165999 | Promoter (<=1kb) | 28126 | ENSMUSG00000025869 |
| H3K27ac | Chchd3 | 9.08 | 9.53 | -0.263259707 | 8.85E-05 | 0.019720478 | Intron | 66075 | ENSMUSG00000053768 |
| H3K27ac | Cds2 | 10.07 | 9.23 | 0.37088334 | 9.63E-05 | 0.021199206 | Promoter (<=1kb) | 110911 | ENSMUSG00000058793 |
| H3K27ac | Hnrnpu | 10.04 | 9.71 | 0.218300713 | 9.94E-05 | 0.021611282 | Promoter (<=1kb) | 51810 | ENSMUSG00000039630 |
| H3K27ac | Gbp7 | 8.68 | 7.83 | 0.368998042 | 0.000100747 | 0.021633562 | Promoter (<=1kb) | 229900 | ENSMUSG00000040253 |
| H3K27ac | Txlng | 9.56 | 9.02 | 0.290581399 | 0.000103154 | 0.021883607 | Promoter (<=1kb) | 353170 | ENSMUSG00000038344 |
| H3K27ac | Gpatch2 | 9.22 | 9.58 | -0.227964081 | 0.000106848 | 0.022322721 | Intron | 67769 | ENSMUSG00000039210 |
| H3K27ac | Pex5 | 9.52 | 9.13 | 0.239593628 | 0.000108447 | 0.022322721 | Promoter (<=1kb) | 19305 | ENSMUSG00000005069 |
| H3K27ac | Desi2 | 8.85 | 9.27 | -0.250754675 | 0.000109027 | 0.022322721 | 5' UTR | 78825 | ENSMUSG00000026502 |
| H3K27ac | Iqsec1 | 12.26 | 12.48 | -0.172809951 | 0.000110792 | 0.022417225 | Promoter (<=1kb) | 232227 | ENSMUSG00000034312 |
| H3K27ac | Atpaf2 | 8.79 | 8.12 | 0.319741457 | 0.000112035 | 0.022417225 | Promoter (<=1kb) | 246782 | ENSMUSG00000042709 |
| H3K27ac | Npr1 | 8.62 | 7.67 | 0.379214165 | 0.000114491 | 0.022651276 | Promoter (<=1kb) | 18160 | ENSMUSG00000027931 |
| H3K27ac | Casp4 | 9.88 | 9.41 | 0.259939018 | 0.000120618 | 0.023421377 | Promoter (<=1kb) | 12363 | ENSMUSG00000033538 |
| H3K27ac | Slc22a15 | 9.66 | 9.99 | -0.219420952 | 0.000122278 | 0.023421377 | Promoter (<=1kb) | 242126 | ENSMUSG00000033147 |
| H3K27ac | Sptssa | 10.02 | 9.62 | 0.240796203 | 0.000122521 | 0.023421377 | Promoter (<=1kb) | 104725 | ENSMUSG00000044408 |
| H3K27ac | Reep4 | 9.57 | 9.09 | 0.26477621 | 0.000124596 | 0.023421377 | Promoter (2-3kb) | 72549 | ENSMUSG00000033589 |
| H3K27ac | Epc2 | 9.47 | 9.87 | -0.23999768 | 0.000125785 | 0.023421377 | Intron | 227867 | ENSMUSG00000069495 |
| H3K27ac | Cast | 10.33 | 9.95 | 0.232923415 | 0.00012737 | 0.023421377 | Promoter (<=1kb) | 12380 | ENSMUSG00000021585 |
| H3K27ac | Phactr2 | 8.81 | 9.30 | -0.2639845 | 0.000128897 | 0.023421377 | Intron | 215789 | ENSMUSG00000062866 |
| H3K27ac | Oprd1 | 9.74 | 10.14 | -0.238004121 | 0.000129025 | 0.023421377 | Distal Intergenic | 18386 | ENSMUSG00000050511 |
| H3K27ac | Jarid2 | 9.13 | 9.56 | -0.246449331 | 0.000132579 | 0.023820887 | Promoter (<=1kb) | 16468 | ENSMUSG00000038518 |
| H3K27ac | Cep19 | 9.15 | 8.63 | 0.270348496 | 0.000140218 | 0.024885344 | Promoter (<=1kb) | 66994 | ENSMUSG00000035790 |
| H3K27ac | Akap8l | 9.47 | 8.97 | 0.258737079 | 0.000141979 | 0.024885344 | Promoter (<=1kb) | 54194 | ENSMUSG00000002625 |
| H3K27ac | Ssbp2 | 9.14 | 9.62 | -0.25947631 | 0.000142743 | 0.024885344 | Exon | 66970 | ENSMUSG00000003992 |
| H3K27ac | Mgat5 | 9.62 | 9.93 | -0.212035169 | 0.000147986 | 0.02554643 | Intron | 107895 | ENSMUSG00000036155 |
| H3K27ac | Mtx1 | 9.63 | 8.65 | 0.36474642 | 0.000151176 | 0.0257936 | Promoter (<=1kb) | 17827 | ENSMUSG00000064068 |
| H3K27ac | Dram1 | 10.57 | 10.11 | 0.254647633 | 0.000152665 | 0.0257936 | Promoter (<=1kb) | 71712 | ENSMUSG00000020057 |
| H3K27ac | Lbr | 10.10 | 9.59 | 0.261424275 | 0.000153812 | 0.0257936 | Promoter (<=1kb) | 98386 | ENSMUSG00000004880 |
| H3K27ac | Dnajb2 | 10.02 | 9.46 | 0.277478543 | 0.000155566 | 0.025841553 | Promoter (<=1kb) | 56812 | ENSMUSG00000026203 |
| H3K27ac | Itpr1 | 9.62 | 9.06 | 0.274614509 | 0.000157328 | 0.02588997 | Promoter (<=1kb) | 16438 | ENSMUSG00000030102 |
| H3K27ac | Ptprm | 10.61 | 10.89 | -0.195244633 | 0.000161511 | 0.026332293 | Intron | 19274 | ENSMUSG00000033278 |

### Table S2 cd.

| Histone modification | SYMBOL | Conc_RSV | Conc_PBS | Fold | p.value | FDR | Genomic annotation | gene ID | ENSEMBL |
|---|---|---|---|---|---|---|---|---|---|
| H3K27ac | Zfp956 | 9.59 | 9.92 | -0.212365344 | 0.000163948 | 0.026484336 | Distal Intergenic | 101197 | ENSMUSG00000045466 |
| H3K27ac | Dhx40 | 8.59 | 7.76 | 0.325955688 | 0.000166863 | 0.026710292 | Distal Intergenic | 67487 | ENSMUSG00000018425 |
| H3K27ac | Samhd1 | 9.77 | 9.22 | 0.266765225 | 0.000192378 | 0.030516987 | Promoter (<=1kb) | 56045 | ENSMUSG00000027639 |
| H3K27ac | Svil | 11.80 | 11.98 | -0.148540966 | 0.000196084 | 0.03082718 | Promoter (<=1kb) | 225115 | ENSMUSG00000024236 |
| H3K27ac | Fgd6 | 9.60 | 8.81 | 0.30992667 | 0.000201404 | 0.031196417 | Promoter (<=1kb) | 13998 | ENSMUSG00000020021 |
| H3K27ac | Magi2 | 8.05 | 8.65 | -0.277080423 | 0.000202165 | 0.031196417 | Intron | 50791 | ENSMUSG00000040003 |
| H3K27ac | Spock1 | 8.53 | 9.01 | -0.253288261 | 0.000203748 | 0.031196417 | Distal Intergenic | 20745 | ENSMUSG00000056222 |
| H3K27ac | Neu3 | 9.06 | 9.48 | -0.231274364 | 0.000209046 | 0.031720034 | 3' UTR | 50877 | ENSMUSG00000035239 |
| H3K27ac | Dlg5 | 8.67 | 9.11 | -0.241070347 | 0.00021077 | 0.031720034 | Promoter (<=1kb) | 71228 | ENSMUSG00000021782 |
| H3K27ac | Id1 | 10.66 | 9.95 | 0.292960075 | 0.000213186 | 0.031811736 | Promoter (<=1kb) | 15901 | ENSMUSG00000042745 |
| H3K27ac | 2700054A10Rik | 9.00 | 9.50 | -0.252142009 | 0.000220732 | 0.032501057 | Promoter (<=1kb) | 72578 | ENSMUSG00000117042 |
| H3K27ac | Vmn1r213 | 8.39 | 8.89 | -0.254716776 | 0.00022232 | 0.032501057 | Distal Intergenic | 171249 | ENSMUSG00000060024 |
| H3K27ac | Lap3 | 9.83 | 8.84 | 0.330150405 | 0.000224494 | 0.032501057 | Promoter (<=1kb) | 66988 | ENSMUSG00000039682 |
| H3K27ac | Patl1 | 10.82 | 10.37 | 0.241313444 | 0.000225715 | 0.032501057 | Promoter (<=1kb) | 225929 | ENSMUSG00000046139 |
| H3K27ac | Usp42 | 9.08 | 7.98 | 0.340733909 | 0.00022805 | 0.032501057 | Promoter (<=1kb) | 76800 | ENSMUSG00000051306 |
| H3K27ac | Dixdc1 | 10.46 | 10.02 | 0.238163408 | 0.000232346 | 0.032501057 | Promoter (<=1kb) | 330938 | ENSMUSG00000032064 |
| H3K27ac | Gsk3b | 12.00 | 12.20 | -0.163044444 | 0.000232583 | 0.032501057 | Promoter (<=1kb) | 56637 | ENSMUSG00000022812 |
| H3K27ac | Gm13056 | 11.19 | 11.00 | 0.159923126 | 0.000233525 | 0.032501057 | Promoter (<=1kb) | 100503810 | ENSMUSG00000085395 |
| H3K27ac | Tanc2 | 9.03 | 9.47 | -0.241238159 | 0.000234418 | 0.032501057 | Intron | 77097 | ENSMUSG00000053580 |
| H3K27ac | Prrc2c | 10.43 | 10.73 | -0.204614064 | 0.000239642 | 0.032946918 | Promoter (<=1kb) | 226562 | ENSMUSG00000040225 |
| H3K27ac | Kdm7a | 8.97 | 9.45 | -0.245033101 | 0.000241683 | 0.032946918 | Exon | 338523 | ENSMUSG00000042599 |
| H3K27ac | Dusp1 | 10.80 | 10.14 | 0.279660001 | 0.000243247 | 0.032946918 | Promoter (<=1kb) | 19252 | ENSMUSG00000024190 |
| H3K27ac | H2-T22 | 9.40 | 8.40 | 0.323957358 | 0.000251019 | 0.033740029 | Promoter (<=1kb) | 15039 | ENSMUSG00000056116 |
| H3K27ac | Inava | 10.16 | 10.41 | -0.182965844 | 0.000264167 | 0.03492163 | Promoter (<=1kb) | 67313 | ENSMUSG00000041605 |
| H3K27ac | Swap70 | 9.65 | 10.07 | -0.232352585 | 0.000265456 | 0.03492163 | Exon | 20947 | ENSMUSG00000031015 |
| H3K27ac | Gm15506 | 9.34 | 8.66 | 0.281394598 | 0.00026576 | 0.03492163 | Promoter (<=1kb) | 100040769 | NA |
| H3K27ac | Irf1 | 8.76 | 7.96 | 0.287285791 | 0.000279905 | 0.036507941 | Promoter (<=1kb) | 16362 | ENSMUSG00000018899 |
| H3K27ac | Ptpa | 9.53 | 9.10 | 0.234623594 | 0.00028399 | 0.036768371 | Promoter (<=1kb) | 110854 | ENSMUSG00000039515 |
| H3K27ac | Kcnip4 | 8.40 | 8.86 | -0.236212086 | 0.000289682 | 0.037231512 | Intron | 80334 | ENSMUSG00000029088 |
| H3K27ac | Nus1 | 9.92 | 9.38 | 0.25204511 | 0.000294669 | 0.037598026 | Promoter (<=1kb) | 52014 | ENSMUSG00000023068 |
| H3K27ac | Hnrnpa0 | 10.25 | 9.60 | 0.266946726 | 0.000304644 | 0.038591222 | Promoter (<=1kb) | 77134 | ENSMUSG00000007836 |
| H3K27ac | Gprc5a | 10.16 | 9.78 | 0.21957316 | 0.000312908 | 0.039111515 | Promoter (<=1kb) | 232431 | ENSMUSG00000046733 |
| H3K27ac | Crebbp | 9.32 | 9.78 | -0.235492855 | 0.000313533 | 0.039111515 | Exon | 12914 | ENSMUSG00000022521 |
| H3K27ac | Fam149b | 8.22 | 7.35 | 0.28743592 | 0.000317337 | 0.039111515 | Promoter (<=1kb) | 105428 | ENSMUSG00000039599 |
| H3K27ac | Pfn2 | 8.11 | 8.63 | -0.247900932 | 0.000317637 | 0.039111515 | Distal Intergenic | 18645 | ENSMUSG00000027805 |
| H3K27ac | Evi5 | 9.15 | 9.52 | -0.219474828 | 0.00032352 | 0.039559291 | Promoter (<=1kb) | 14020 | ENSMUSG00000011831 |
| H3K27ac | Zbtb7b | 10.81 | 10.34 | 0.237466218 | 0.000326496 | 0.039647858 | Promoter (<=1kb) | 22724 | ENSMUSG00000028042 |
| H3K27ac | Irgm2 | 8.82 | 7.98 | 0.282203892 | 0.000339383 | 0.04093046 | Promoter (<=1kb) | 54396 | ENSMUSG00000069874 |
| H3K27ac | Gmpr | 9.17 | 9.51 | -0.209567553 | 0.000351437 | 0.041952647 | 3' UTR | 66355 | ENSMUSG00000000253 |
| H3K27ac | Eif2ak3 | 10.38 | 9.71 | 0.262763887 | 0.000352623 | 0.041952647 | Promoter (<=1kb) | 13666 | ENSMUSG00000031668 |
| H3K27ac | Paip2b | 9.73 | 9.08 | 0.25935961 | 0.000366454 | 0.043305569 | Promoter (<=1kb) | 232164 | ENSMUSG00000045896 |
| H3K27ac | Zfp708 | 9.68 | 9.01 | 0.2597219 | 0.000371279 | 0.04356128 | Promoter (<=1kb) | 432769 | ENSMUSG00000058883 |
| H3K27ac | Stard13 | 11.50 | 11.71 | -0.159845264 | 0.000373566 | 0.04356128 | Promoter (<=1kb) | 243362 | ENSMUSG00000016128 |
| H3K27ac | Sned1 | 8.17 | 8.68 | -0.240615379 | 0.000383519 | 0.044236221 | Exon | 208777 | ENSMUSG00000047793 |
| H3K27ac | Tapt1 | 11.93 | 12.15 | -0.164727412 | 0.000384379 | 0.044236221 | Promoter (<=1kb) | 231225 | ENSMUSG00000046985 |
| H3K27ac | Abl1 | 9.45 | 9.93 | -0.235068046 | 0.000401019 | 0.045712193 | Exon | 11350 | ENSMUSG00000026842 |
| H3K27ac | Txndc2 | 8.47 | 8.91 | -0.224760024 | 0.000402396 | 0.045712193 | Intron | 213272 | ENSMUSG00000050612 |
| H3K27ac | Jdp2 | 10.03 | 9.72 | 0.200814476 | 0.00040612 | 0.045839532 | Promoter (<=1kb) | 81703 | ENSMUSG00000034271 |
| H3K27ac | Ubc | 11.66 | 11.33 | 0.203890834 | 0.000411257 | 0.046123602 | Promoter (<=1kb) | 22190 | ENSMUSG00000008348 |
| H3K27ac | Vgll4 | 10.84 | 10.51 | 0.204714434 | 0.000415077 | 0.046257476 | Promoter (<=1kb) | 232334 | ENSMUSG00000030315 |
| H3K27ac | Zfp239 | 9.53 | 8.76 | 0.260634687 | 0.000423292 | 0.046876266 | Distal Intergenic | 22685 | ENSMUSG00000042097 |
| H3K27ac | Bmp3 | 9.45 | 9.79 | -0.202544634 | 0.00042837 | 0.047142165 | Distal Intergenic | 110075 | ENSMUSG00000029335 |
| H3K27ac | Ift172 | 9.14 | 8.32 | 0.262642001 | 0.000441306 | 0.047826639 | Promoter (<=1kb) | 67661 | ENSMUSG00000038564 |
| H3K27ac | Foxp2 | 9.59 | 9.98 | -0.214929525 | 0.000442487 | 0.047826639 | Promoter (<=1kb) | 114142 | ENSMUSG00000029563 |
| H3K27ac | Brpf3 | 9.07 | 7.72 | 0.268982481 | 0.000443473 | 0.047826639 | Promoter (<=1kb) | 268936 | ENSMUSG00000063952 |
| H3K27ac | Tmem253 | 10.62 | 10.14 | 0.230041043 | 0.000446866 | 0.047826639 | Promoter (<=1kb) | 619301 | ENSMUSG00000072571 |
| H3K27ac | Parp9 | 8.14 | 7.22 | 0.266491364 | 0.000448171 | 0.047826639 | Distal Intergenic | 80285 | ENSMUSG00000022906 |
| H3K27ac | Psmb9 | 8.51 | 7.47 | 0.270026385 | 0.000457337 | 0.048510805 | Promoter (<=1kb) | 16912 | ENSMUSG00000096727 |
| H3K27ac | Rpusd3 | 8.30 | 7.50 | 0.257607425 | 0.000468933 | 0.049442992 | Promoter (<=1kb) | 101122 | ENSMUSG00000051169 |
| H3K27ac | Firre | 9.02 | 8.48 | 0.236036663 | 0.000475632 | 0.049850784 | Exon | 103012 | ENSMUSG00000085396 |
